# Increased versatility and convenience: advances and strategy optimization of ReMOT-mediated genetic modification in insects

**DOI:** 10.1101/2024.12.24.629949

**Authors:** Xia Ling, Cao Zhou, Jun-feng Hong, Yan-ping Jiang, Ling-yi Li, Si-yi Wang, Xin-yuan Xie, Qi-li Zou, Xiao-ling Yang, Kai Xiang, Jin Ma, Liang Qiao, Chen Bin, Wei Sun

## Abstract

Genetic modification via gene editing has become a widely adopted and demonstrably effective method in functional gene research within entomology. However, the optimal efficiency and simplicity of delivering exogenous guide RNA-clustered regularly interspaced short palindromic repeats-associated protein 9 (gRNA-Cas9) complexes into target tissues are crucial for successful gene editing. The Receptor-Mediated Ovary Transduction of Cargo (ReMOT) strategy, which simplifies the delivery process, target site selection, technical requirements, and delivery cost compared with embryonic microinjection, enabling efficient editing at the germline level, is gaining increasing attention. Although the feasibility and advantages of this technique have been demonstrated in various insect species, further optimization of operational details and the overcoming of further bottlenecks are still required. This review focuses on advances in developing ReMOT as a valuable technology, exploring its applicability, rationale for selecting the ovary as a delivery target site, factors influencing its efficiency, and recommendations for improvement. The versatility and effectiveness of ReMOT make it a promising method for researchers looking to make precise genetic modifications with greater ease and efficiency.

## Introduction

Insects represent the most diverse group of animals on Earth, making them an excellent material for studying species diversity and differentiation, the evolution of adaptive traits, metamorphosis, and body pattern development, among others (Scudder 2009, Tihelka et al. 2021, Truman 2019). Furthermore, research on applied issues, such as the domestication and breeding of economically important insects, pest control, and the disease transmission mechanisms of insect vectors, is directly related to impacts on human society, including food security and the economy (Belluco et al. 2023, Hill 1997, Paganizza 2017). The key to addressing these scientific issues lies in deciphering the genetic basis underlying the biological phenomena of interest. Clustered regularly interspaced short palindromic repeats (CRISPR)/CRISPR-associated protein 9 (Cas9)-mediated gene editing, which utilizes the specific recognition between RNA and DNA to guide Cas9 to the target genetic locus, has become a powerful tool for DNA functional analysis because of its versatility and the advantage of not requiring the redesign of nucleases for different target genes (Ansori et al. 2023). Currently, this approach is used to analyze the molecular basis of numerous biological questions and to elucidate the molecular mechanisms of causative genes or genetic elements in insects as well as other organism groups.

Simple and efficient methodologies for delivering exogenous nucleic acid-protein complexes are vital for the achievement of effective and stable genetic modification *in vivo*. For example, the ligand region at the N terminus of the *Drosophila melanogaster* Yolk protein 1 (Yp1) efficiently recognized and bound by oocyte membrane receptors; by incrementally truncating the N terminus of Yp1, Chaverra-Rodriguez et al. identified a 41-amino acid-guiding peptide, P2C, which was found to direct Cas9 to be specifically endocytosed into the ovaries of female mosquitoes (Chaverra-Rodriguez et al. 2018). This discovery led to the development of the Receptor-Mediated Ovary Transduction of Cargo (ReMOT) method, which enables the targeted delivery of gRNA-Cas9 complex into the ovaries of adult female insects, thereby facilitating germline-level genome editing of oocytes or embryos (Chaverra-Rodriguez et al. 2018). Since then, gene-editing research based on the principles of the ReMOT strategy has expanded rapidly in insects and other animals. In response to this emerging gene-editing strategy, we review recent advances in the main use of ReMOT in arthropods, with a focus on insects, and discuss its applicability and advantages, the criteria for selecting the delivery target site, factors affecting the efficiency of ReMOT implementation, and recommendations for improvement. Additionally, we explore the potential power and extensions of genetic modification techniques in insects based on the principles of ReMOT.

### 1. Species applicability and technical advantages of ReMOT delivery

The strategy of delivering exogenous nucleic acid-protein complexes into maternal organisms has been applied to over 30 arthropod species, with a total of 80 documented cases (Figure 1, Table S1). Among these, 42.5% (34/80) of deliveries were achieved using the ReMOT technique (injection of Cas9 with a ligand) sensu stricto, while 57.5% (44/80) were performed via injection of Cas9 without a ligand (e.g., Direct Parental (DIPA) Cas9, Branched Amphiphilic Peptide Capsules (BAPC) Cas9, and Synergistic CRISPR-Cas Systems (SYNCAS), etc.) (Figure 1, Table S1). In terms of genetic modifications, the targeted gene types include those involved in eye pigment formation, cuticular coloration, physiological metabolism, and development modes related to body plans (Chaverra-Rodriguez et al. 2018, Cohen et al. 2023, Iwaizumi et al. 2021, Terradas et al. 2022, Yu et al. 2023, Zhang et al. 2023, Dermauw et al. 2020, De Rouck et al. 2024, İnak et al. 2024b, Mocchetti et al. 2025, Zhang et al. 2024, Shirai et al. 2022, Shirai and Daimon 2020, Chaverra-Rodriguez et al. 2020, Dalla Benetta et al. 2020, Lima et al. 2024, Sharma et al. 2022, Kondo et al. 2024, Liu et al. 2024, İnak et al. 2024a, Miller 2021, Hunter et al. 2018). In vertebrates such as zebrafish, fluorescent proteins fused with Vtg-ligand peptides synthesized in the female parent’s liver can also be selectively absorbed by oocytes through receptor-mediated endocytosis (Iwaizumi et al. 2021). These findings suggest that the strategy of delivering RNPs via maternal injection has relatively broad applicability. However, the underlying mechanisms for methods such as DIPA, BAPC Cas9 and SYNCAS remain to be elucidated. These mechanisms have been hypothesized to rely on the random diffusion of delivery complexes and nonspecific cellular uptake. Consequently, the implementation of DIPA, BAPC Cas9 and SYNCAS methods requires careful consideration of their applicability to the intended species, the consequences of delivery complex dosage on female parents and the potential for Non targeted effects on non-target tissues. In contrast, ReMOT uses a targeted delivery mechanism involving the recognition and binding of specific vitellogenin receptors and vitellogenin ligand regions (Chen et al. 2023, Stein et al. 2022, Terradas et al. 2023, Chen and Palli 2024), ensuring precise delivery of exogenous RNP complexes to germ cells. Theoretically, the delivery of exogenous nucleic acids should be achievable, on the basis of the principles of ReMOT in any species in which vitellogenesis occurs, including arthropods.

**Figure 1.**
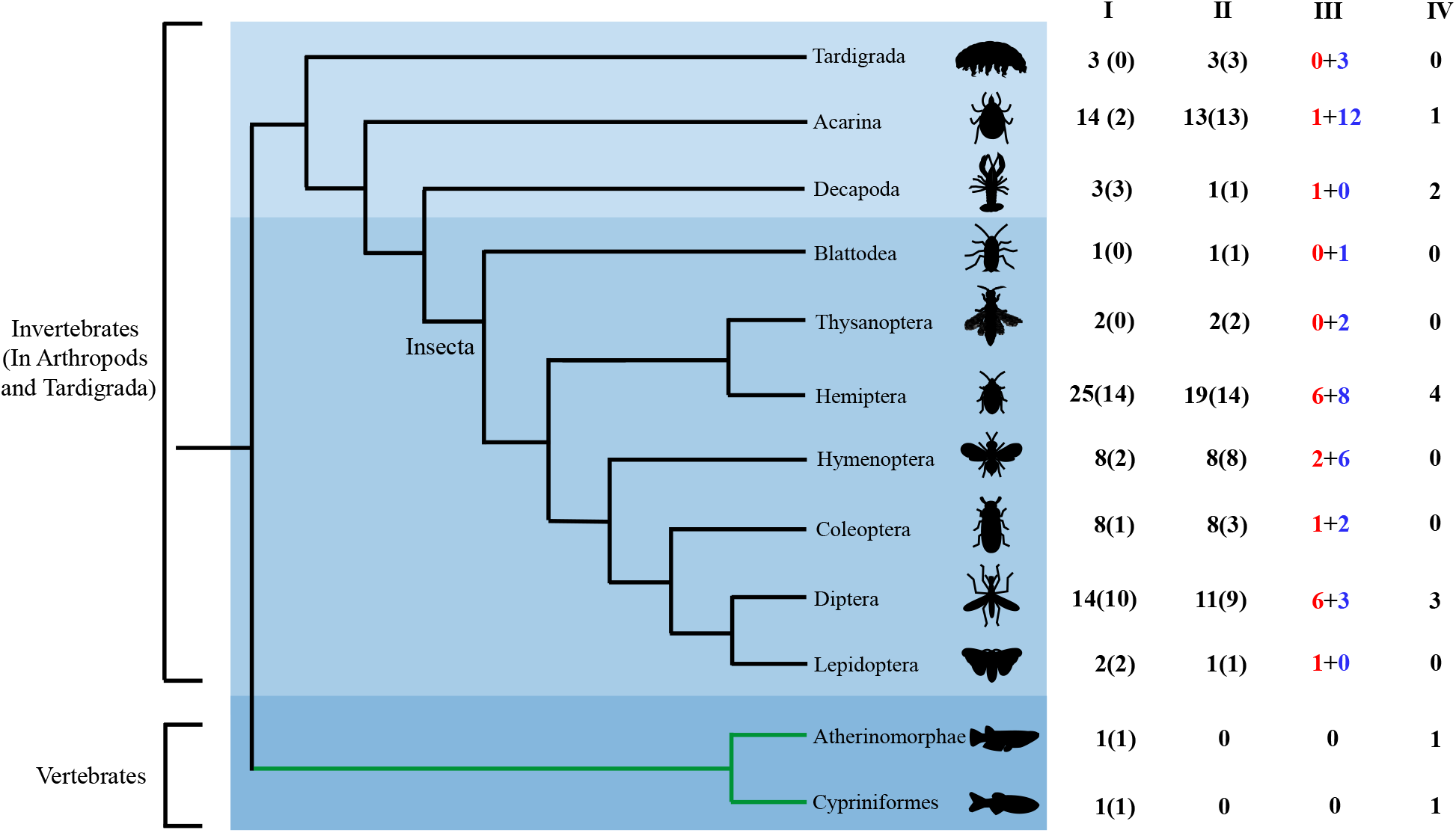
Examples of studies mediated by maternal injection for delivery to female germ cells in insects and other animals. Column I represents the cases of delivery mediated by maternal injection the number of cases using peptide ligands is shown in parentheses. Column II represents the cases of delivery involving genetic modification (including gene editing and transient gene silencing); the number of successful genetic modifications indicated in parentheses. Column III represents the cases in which genetic modification was successfully achieved; the numbers of cases that used ovarian peptide ligands (in red characters) and those that did not use peptide ligands (in blue characters) are shown. Column IV represents the cases of maternal injection-mediated delivery that used peptide ligands and achieved targeted delivery to the ovaries, but did not involve or contribute to particular genetic modifications.

**Figure 2.**
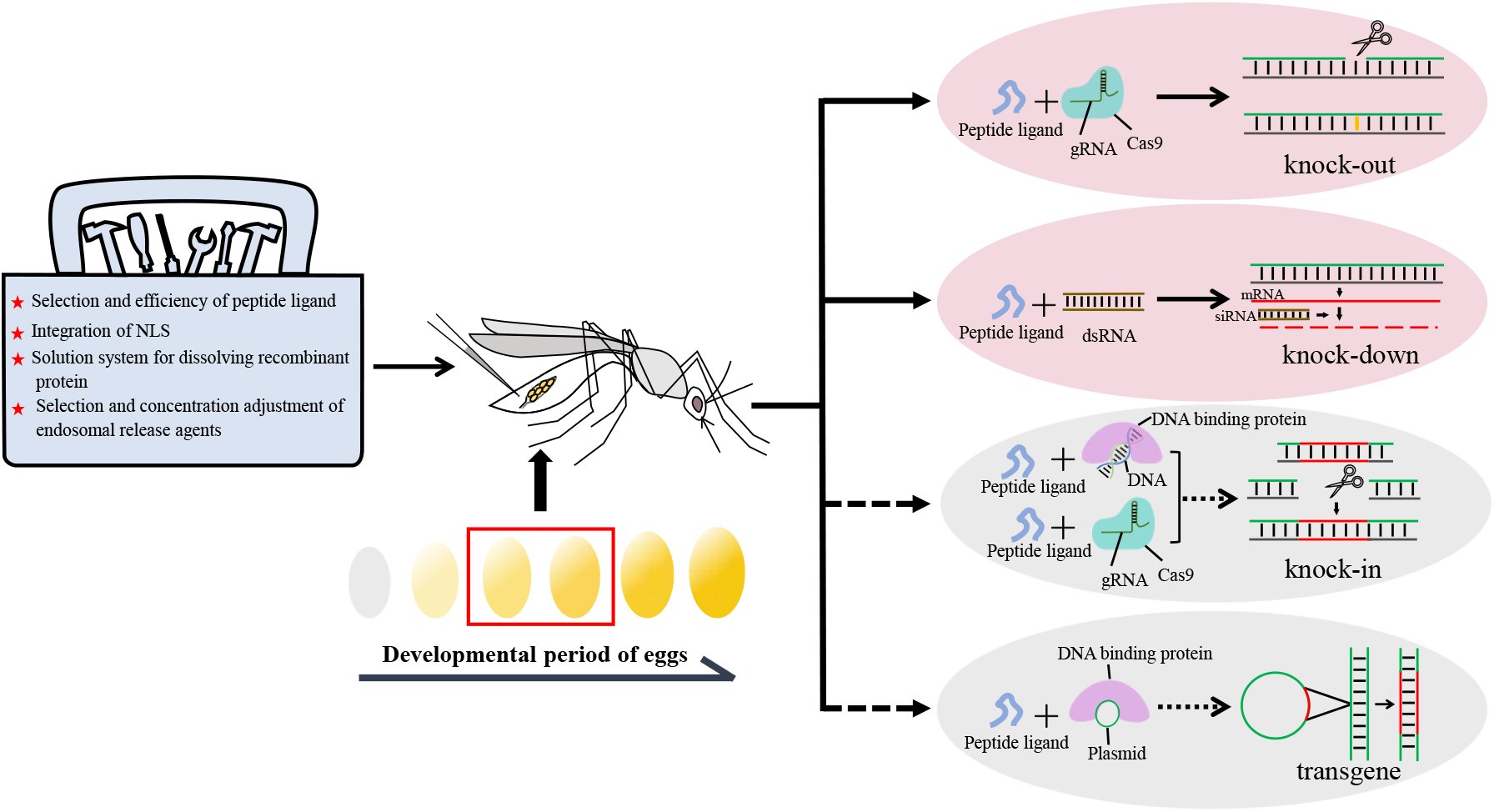
Workflow for genetic modification based on ReMOT delivery in animals (represented by insects). Currently, ReMOT can be used to deliver DNA molecules into female germ cells. In the future, ReMOT could be used to achieve more diverse genetic modifications, such as gene knock-in and transgenesis, in addition to already-realized gene knockout and knockdown. The elliptical circle represents the eggs. Vitellogenin (yellow) gradually accumulates during the eggs’ developmental period. In mosquitoes, for example, injection of RNP complexes into the maternal body is usually performed at the vitellogenesis stage (indicated within the red rectangle). The red stars represent the four key aspects that must be considered for optimizing ReMOT technology. Dashed arrows represent the forms of genetic modifications that can be achieved through ReMOT technology.

For germline mutagenesis, although embryonic microinjection is the most common manipulation for performing CRISPR/Cas9-mediated gene editing, operational difficulties during the injection process and the survival rate of individuals post injection significantly affect the resulting delivery and editing efficiency (Criscione et al. 2015). Given that, in insects, particularly for non-model species, the recipients of ReMOT delivery are adults, the technical requirement threshold is significantly reduced compared with embryonic injection, which makes it easier to master the approach. For example, in species such as *Diaphorina citri*, the survival rate of adults after ReMOT injection ranges from 60% to 80%, and most surviving individuals exhibit normal egg-laying ability (Chaverra-Rodriguez et al. 2023). Consequently, the number of viable samples available for candidate screening following a single injection is increased.

In terms of the versatility of the technology used for embryo microinjection, the parameters suitable for one insect species (such as needle size, injection stage, or post-injection incubation conditions) might not be appropriate for others, especially non-model insects (Carballar-Lejarazu et al. 2021, Lobo et al. 2006, Ringrose 2009). This makes the trial-and-error process of embryo microinjection very complex and cumbersome. Furthermore, the cost of embryo microinjection equipment is high, with expensive consumables and maintenance costs. Moreover, in viviparous insects such as tsetse flies and aphids, performing zygotic gene editing via embryonic injection is impossible (Benoit et al. 2015, De Vooght et al. 2018, Jamison et al. 2018). However, the fundamental principles and distinctive characteristics of ReMOT suggest the potential to achieve germline-level genetic modification even in these species. Therefore, the ReMOT delivery system demonstrates significant advantages in terms of user-friendly and practical and easy to use.

### 2. Selection criteria for ovarian tissue as the target site of ReMOT delivery

The ReMOT technique focus on targeted delivery to oocytes. Theoretically, delivering RNP complexes to both male and female germ cells could achieve germline-level editing. However, there are several practical challenges to targeting sperm for delivery. On the one hand, efficient peptide ligands for targeting sperm still need to be developed. On the other hand, even if efficacious male germline cell peptide ligands were identified, it might be necessary to deliver extremely high concentrations or large doses of RNP complexes to target most of the sperm to achieve effective delivery and editing, given the abundance of sperm in the testes compared with oocytes in ovaries. This is in addition to any potential toxic or lethal effects. Even if a few sperm received sufficient doses of the RNP complexes and undergo editing, the probability of these few ‘lucky’ sperm fertilizing eggs would still be low. These factors could result in a very high proportion of unedited zygotes. Consequently, the use of guide peptides targeting oocytes is a more suitable approach and holds greater potential for further exploration.

### 3. Species specificity of peptide ligands correlated with delivery efficiency

Ovary delivery-mediated gene editing could also be achieved by directly injecting Cas9 (DIPA Cas9) without guide peptides in *Tribolium castaneum, Blattella germanica and Aedes aegypti* (Shirai et al. 2022, Shirai et al. 2023). The insects in which this approach can be used successfully are characterized by highly efficient RNAi delivery, although the specific correlation is not yet clear. A screen of previous cases involving ovarian delivery of RNP complexes indicates that, in most insects, the complexes require the peptide ligand to efficiently enter germ cells and complete editing events. In vertebrates, such as the Japanese rice fish *Oryzias latipes*, the vitellogenin (Vg) signal region has also been reported to mediate the delivery of exogenous proteins into the oocytes (Murakami et al. 2019). During the vitellogenesis stage of insect oogenesis, yolk proteins and other nutrients are absorbed and stored in oocytes through receptor-mediated endocytosis (Izumi et al. 1994, Raikhel and Dhadialla 1992, Sappington and Raikhel 1998, Snigirevskaya et al. 1997). The ligand region of the yolk protein, also designated as the peptide ligand, binds to receptors on the oocyte membrane, enabling Vg to enter the oocyte. This short ligand region can be identified by sequential truncation of the amino acid sequence of major intra-oocyte nutrients at the cellular or individual level.

The peptide ligand from yolk protein 1 of *Drosophila melanogaster* (‘P2C’) or Vg from the *Bemisia tabaci* (‘BTKV’) has been identified used in previous gene editing studies (Heu et al. 2020, Chaverra-Rodriguez et al. 2018). These peptides are fused with Cas9 to direct the absorption of RNP by oocytes. However, within the same species, the use of different types of peptide ligand can result in significant variation in both the delivery and final editing efficiencies(Heu et al. 2020, Yu et al. 2023). This suggests that the compatibility of ligand and receptor amino acid sequences is crucial for their interaction.

Earlier studies have confirmed that the β-sheet region at the N-terminus of yolk proteins from various oviparous animals contain animo acid sites that interact with the Vg receptor, which is considered crucial for receptor recognition of the ligand and directed uptake into the oocytes (Roth et al. 2013). Consequently, the regions potentially recognized and bound by the VgR might potentially be inferred by examining the N-terminal sequences of existing yolk proteins. A comparison of potential peptide ligands in N-terminus region among various insect yolk proteins revealed that, although P2C from the *D. melanogaster* Yp1 matched the N-terminal region of Yp1 homologs in other Brachycera (Diptera), there was substantial sequence diversity within this suborder (Table 1, Table S2 and Figure S1). Concurrently, although the ligand peptide BTKV (from the N-terminal region of *Bemisia tabac*i’s Vg) as a seed sequence matches the N-terminal region of Vg in non-dipteran insects, the predicted ligand peptide regions still exhibit diversity among orders (Table 1, Table S2 and Figure S1). In arthropods outside the *Insecta*, such as *Parasitiformes* mites and *Decapoda* crustaceans, as well as vertebrate groups, such as *Atherinomorphae* and *Cypriniformes*, the predicted peptide ligand motifs in Vg, which bind to the Vg receptor (VgR), also exhibit order-specific characteristics (Table 2, Table S2 and Figure S1). We hypothesize that, if the peptide ligand from other species significantly differs from those of the target species, this would weaken the binding of the peptide ligand to the corresponding membrane receptor, ultimately affecting the efficiency of delivery into the oocytes. Therefore, the prediction and screening of peptide ligands are crucial for efficient delivery. It is suggested that peptide ligand sequences should be selected from the target species or the conserved peptide segment within the taxonomic order to which the species belongs. Additionally, in the case of Lepidoptera, oocytes absorb 30k proteins during development as nutrients, in addition to Vg (Zhang et al. 2012). These proteins enter oocytes via cell-penetrating peptides (CPPs), rather than relying on specific receptor-mediated transportation. Current research also explores the use of CPPs fused with Cas9 to mediate RNP delivery to cells (Darif et al. 2023, Falato et al. 2022). However, CPPs do not exhibit specificity in terms of the types of cells they enter. If CPPs are used to guide Cas9, a substantial amount of the fusion protein must be injected into the female to ensure that sufficient RNP enters the germ cells to achieve gene editing. This increases the cost, protein complexity, and difficulty associated with their synthesis. Furthermore, the excessive accumulation of exogenous proteins could lead to cytotoxicity.

**Table 1.**
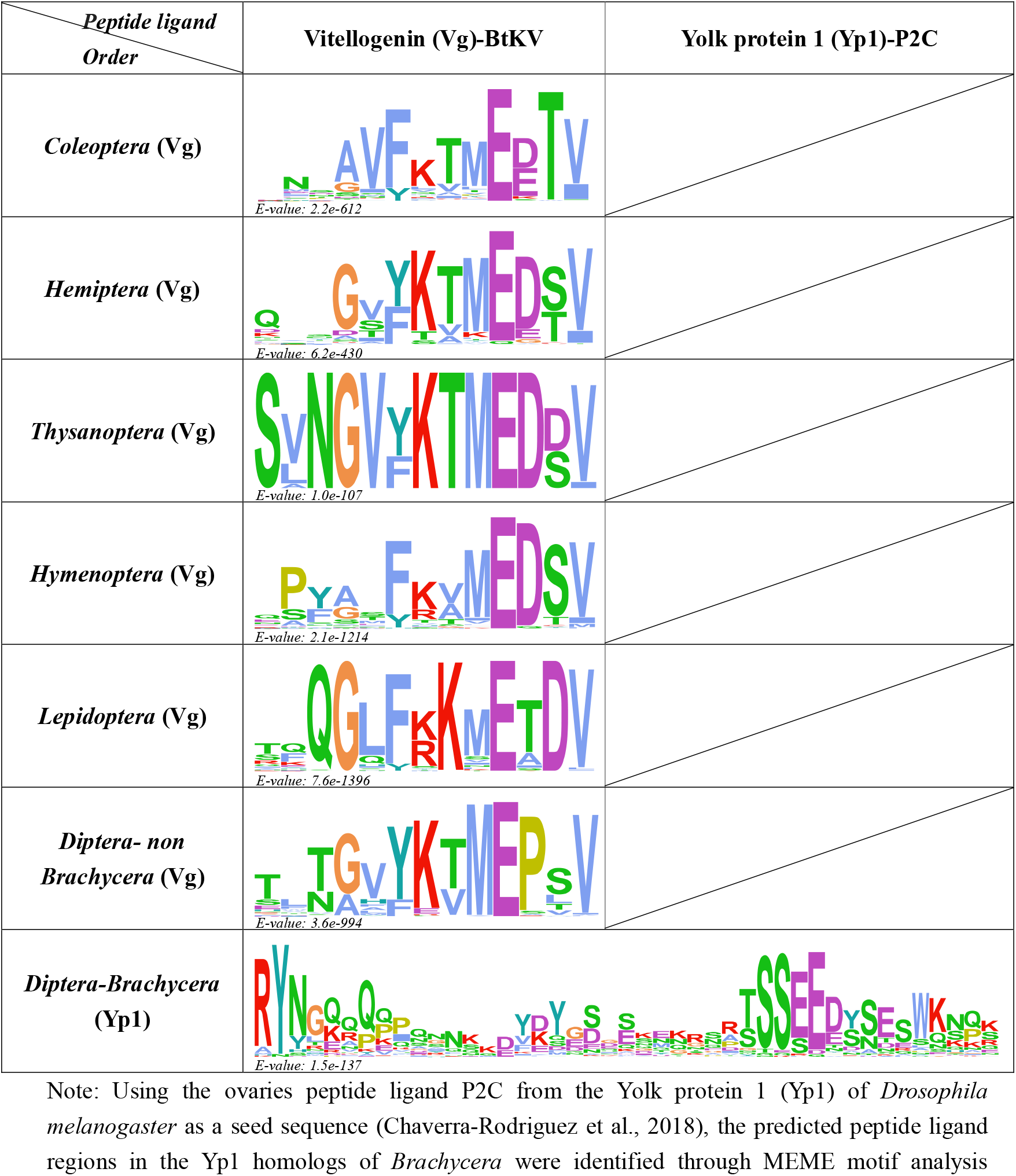
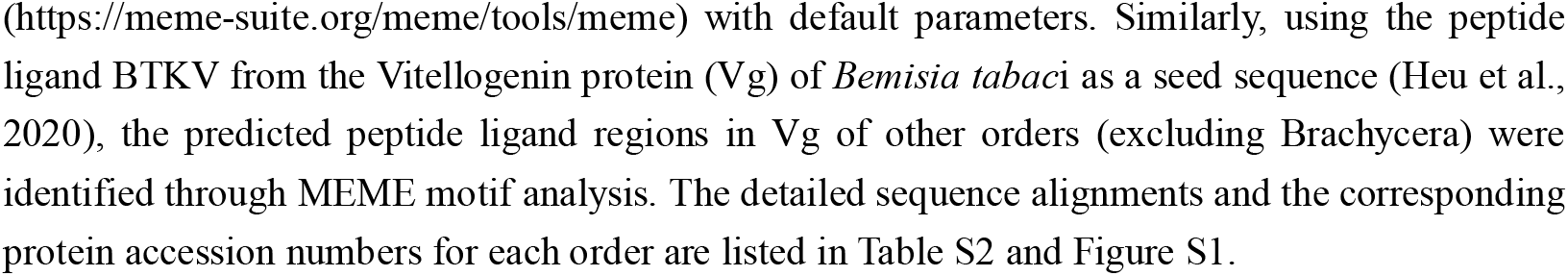
Predicted ovarian peptide ligand regions in selected insect orders.

**Table 2.**
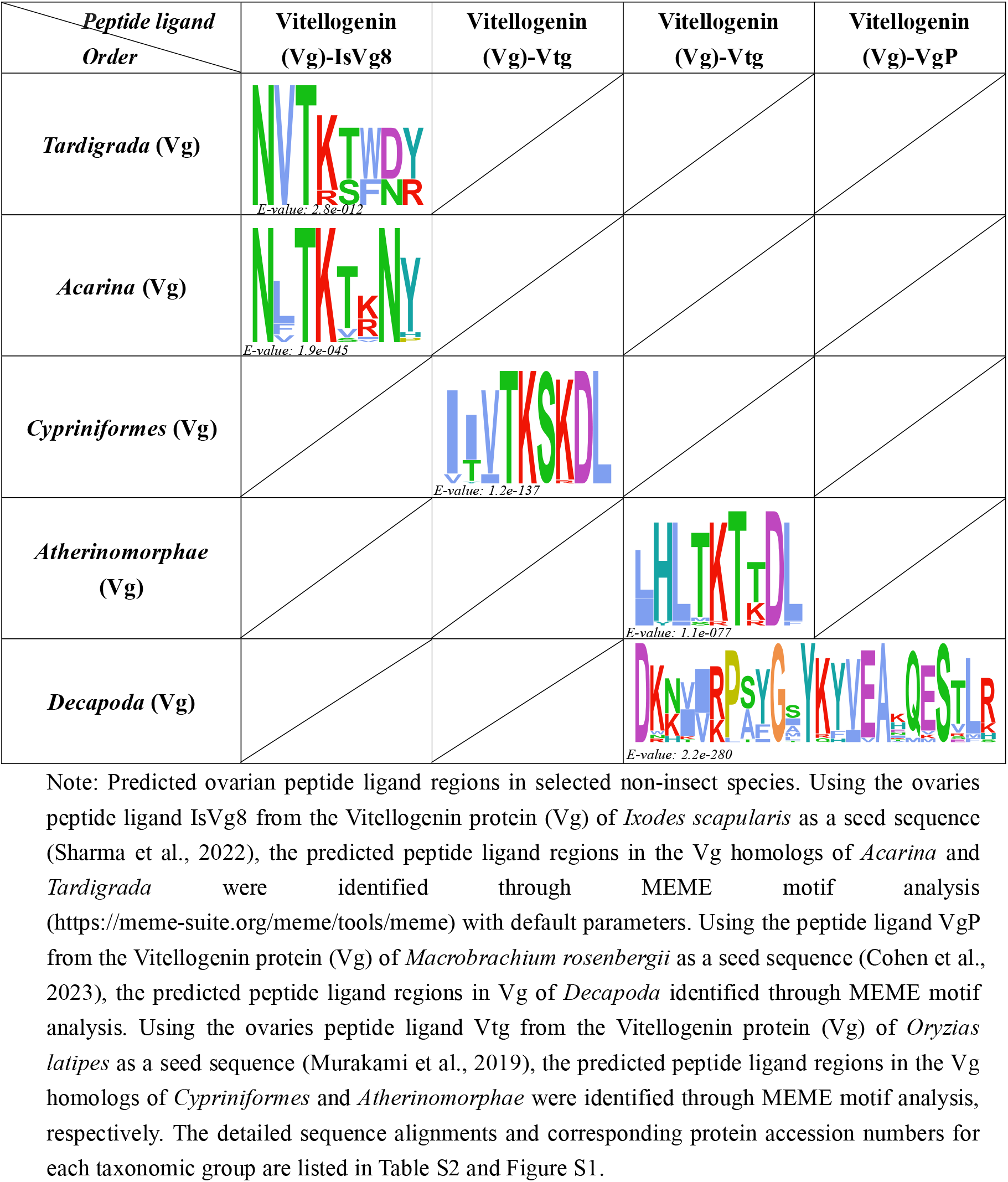
Predicted ovarian peptide ligand regions in selected non-insect species.

| *Peptide ligand Order* | Vitellogenin (Vg)-IsVg8 | Vitellogenin (Vg)-Vtg | Vitellogenin (Vg)-Vtg | Vitellogenin (Vg)-VgP |
| --- | --- | --- | --- | --- |
| *Tardigrada* (Vg) | E-value: 2.8e-012 |  |  |  |
| *Acarina* (Vg) | E-value: 1.9e-045 |  |  |  |
| *Cypriniformes* (Vg) |  | E-value: 1.2e-137 |  |  |
| *Atherinomorphae* (Vg) |  |  | E-value: 1.1e-077 |  |
| *Decapoda* (Vg) |  |  |  | E-value: 2.2e-280 |

Note: Predicted ovarian peptide ligand regions in selected non-insect species. Using the ovaries peptide ligand IsVg8 from the Vitellogenin protein (Vg) of *Ixodes scapularis* as a seed sequence (Sharma et al., 2022), the predicted peptide ligand regions in the Vg homologs of *Acarina* and *Tardigrada* were identified through MEME motif analysis (https://meme-suite.org/meme/tools/meme) with default parameters. Using the peptide ligand VgP from the Vitellogenin protein (Vg) of *Macrobrachium rosenbergii* as a seed sequence (Cohen et al., 2023), the predicted peptide ligand regions in Vg of *Decapoda* identified through MEME motif analysis. Using the ovaries peptide ligand Vtg from the Vitellogenin protein (Vg) of *Oryzias latipes* as a seed sequence (Murakami et al., 2019), the predicted peptide ligand regions in the Vg homologs of *Cypriniformes* and *Atherinomorphae* were identified through MEME motif analysis, respectively. The detailed sequence alignments and corresponding protein accession numbers for each taxonomic group are listed in Table S2 and Figure S1.

### 4. Optimization strategies for ReMOT implementation

ReMOT delivery methods utilize recombinant proteins fused with the peptide ligand and Cas9, developed by researchers, except for direct parental (DIPA) delivery, which uses commercial Cas9 protein without a peptide ligand (Dermauw et al. 2020, Shirai et al. 2022, Zhang et al. 2024, Matsuda et al. 2024, Terradas et al. 2023). During the construction of protein expression plasmids and purification of recombinant proteins, it is vital to consider several crucial components and modifications: (a) the inclusion of the nuclear localization signal (NLS), such as SV40 NLS or nucleoplasmin NLS, which is essential for the nuclear entry of the RNP complex, although the necessity of linkers has not yet been validated; (b) visual markers, such as EGFP or DsRed, to directly indicate the delivery efficiency of the peptide ligand. However, studies show that larger molecular weights of fused Cas9 protein can reduce the editing efficiency (Chaverra-Rodriguez et al. 2018). Consequently, it is recommended that, once the delivery efficiency of the peptide ligand has been confirmed, there is no need to integrate the coding sequence of the reporter gene into the peptide ligand (Yang et al. 2024); (c) it is also necessary to consider the yield and desalting of recombinant proteins, because these are crucial factors that affect the injection dosage, the chemical environment for *in vitro* incubation, and any toxicity effects on the injected organism; (d) it is recommended that the recombinant protein fused with Cas9 and peptide ligand should be subjected to controlled *in vitro* split activity assays (including comparisons with commercial Cas9 proteins) to assess the activity and ensure editing efficiency *in vivo* (Yang et al. 2024).

Adding saponin or chloroquine to the delivery RNP complex can promote endosomal escape. In mosquitoes, the incorporation of saponin resulted in higher gene-editing efficiency (Macias et al. 2020). Furthermore, the injection of varying types and concentrations of endosomal escape agent markedly influenced the survival rates of the species under examination (Li et al. 2021, Macias et al. 2020, Yang et al. 2024, Yu et al. 2023). Therefore, it is necessary to screen for endosomal escape agents that facilitate editing and that show low toxicity or are nontoxic, to identify those that are suitable for foreign nucleic acid delivery via ReMOT for use with specific insects. Moreover, when determining the injection dosage, different concentrations of endosomal escape agents should be tested for their effects on adult mortality and larval hatching rates to establish an appropriate range of escape agent concentrations that balance editing efficiency with the number of surviving adults.

In the mosquito *A. aegypti*, the planthopper *Sogatella furcifera*, and the flour beetle *Tribolium castaneum*, the editing efficiency of gRNA-Cas9 complexes varies with different ovarian developmental stages (Chaverra-Rodriguez et al. 2018, Macias et al. 2020, Shirai et al. 2023, Shirai et al. 2022). Based on these results, it was found that delivering RNP complexes during early to mid-stages or slightly earlier in the mid-stage of ovary development led to higher editing and incorporation efficiency in the offspring compared with early and late stages of ovary development. We speculate that the oocyte has the strongest capacity to endocytose external nutrients during vitellogenesis. At this time, the deposition of nutrients within the egg is not yet sufficiently dense. Consequently, the RNP complexes fused with peptide ligands are likely to be endocytosed extensively as ‘mimicked’ nutrients. Additionally, there might be more spatial opportunities for contact with the nucleus. Thus, before conducting ReMOT delivery in the target insect species, it is essential to analyze the accumulation pattern of nutritional substances during different stages of oogenesis. This can be achieved through a range of techniques, including dissection of the internal reproductive glands, ovarian sectioning, immunostaining, and others. Additionally, investigating the temporal and spatial expression patterns of *Vg* and other yolk protein-coding genes in female adults will assist in identifying the peak expression stages. The results of these studies will collectively contribute to the determination of the optimal injection time.

### 5. Summary and perspective

The efficacy of ReMOT-mediated gene editing has been demonstrated in a range of species, with particular success observed in insects. The universal applicability of its fundamental principles and easy accessibility for achieving technical thresholds, and its balance between investment and efficiency make it particularly attractive. Furthermore, in certain industrial applications, such as the genetic improvement of silkworm strains with egg diapause, ReMOT delivery can avoid the extremely high mortality rate caused by acid treatment required to break diapause, followed by embryonic injection. In pest genetic control, especially for those species that are difficult to manipulate at the embryonic stage, ReMOT-mediated gene editing can facilitate a quicker and more straightforward survey of target genes. Currently, genetic modifications utilizing ReMOT primarily concentrate on gene-knockout analysis through the delivery of gRNA-Cas9 complexes containing oocyte peptide ligand. In theory, <u>if</u> recombinant proteins with peptide ligands facilitate the targeted delivery of DNA molecules to germ cells, this approach should enable more diverse and complex molecular genetic operations, such as gene knock-in and transgenesis, with less stringent technical requirements. Recent research on extracellular contractile injection systems (eCISs) suggest that the feasibility of fusing tissue or organ-specific peptide ligands with viral or bacterial capsid proteins to encapsulate DNA molecules, such as gene knock-in templates and genetic modification plasmids(Kreitz et al. 2023). This strategy could enable the targeted ‘infection’ of germ cells, facilitating the delivery of exogenous DNA to achieve transient or germline-level genetic modifications. Additionally, recombinant proteins fused with peptide ligands and the DNA-binding domain of the Gal4 transcription factor have been shown to carry transgenic plasmids containing specific UAS motif sequences into mosquito ovarian tissues, providing new insights into exogenous DNA delivery technology built upon ReMOT (Yang et al. 2023). This review of the progress in ReMOT and its suggested use for the optimization and expansion of the gene-editing approach undoubtedly shows the broader applications of this technology in a wider range of animal groups, including insects, and also highlights the potential for more diverse and refined genetic modifications.

## Acknowledgement

This work was supported by the National Natural Science Foundation of China (Nos. 31772527, 31872262), the Scientific and Technological Research Program of Chongqing Municipal Education Commission (Nos. KJZD-K202200507; KJQN202200533); the Venture and Innovation Support Program for Chongqing Overseas Returnees (No. cx2022052), Chongqing “Express” Science and Research Program for PhD (No. CSTB2022BSXM-JCX0067), the Graduate Research and Innovation Foundation of Chongqing under Grant (Nos. CYS23399, CYS240374) and College Students’ Innovation and Entrepreneurship Training Plan Program (No. 202410637012).

## SUPPORTING INFORMATION

**Table S1**. Summary of cases involving the delivery of exogenous nucleic acid-protein complexes into the maternal body in arthropods and other animals. Orange shading represents delivery mediated by the ovarian peptide ligand, whereas green shading represents other delivery methods. “GEF” denotes G_0_ gene-editing frequency, “EEF” denotes effort efficiency, and an asterisk (*) indicates the editing efficiency calculated from the data provided in the literature. A hashtag (#) indicates that only the number of genetic modified individuals is provided, but the number of offspring is not provided.

**Table S2**. Organisms used for analysis of predicted ovarian peptide ligand regions.

**Figure S1**. Detailed alignments of predicted ovarian peptide ligands across various insect groups and other animal groups.

## Notes

### Competing Interest Statement

The authors have declared no competing interest.

