## Supplementary material for "Increased versatility and convenience: advances and strategy optimization of ReMOT-mediated genetic modification in insects": Table S1, Table S2, Figure S1: 20241224 Figure S1.pdf

#### *Coleoptera* (Vg)

β-sheet

predicted peptides

```
XP_017781017 PQ-EG-SNNAVFKTMEDTVTGKYETLYDITPLPEYILQ----SQPWLVPMPQ 46
XP_017778650 PE-AD-SDSAVFKTMEETVTGKYETLYEISPLPEYVLQ----SQPWLVPMPE 46
KAF2882317   PE-QG--PYGVYKTMEETVTGKYQTLYELSPLPEYYYE----NQPWEVPYPE 45
XP_019880305 PN--N-NNNGVYKTMEDTVNGECETLYDIHPLPEYILE----SKPWLAPQQQ 45
CAG9759827   PN-GE-RMDGVFKTMEETVTGKTETLYDIHPLPEYLLQ----SEPWLAPQDS 46
XP_971398    TQ-ED-SNTAVFKTMEETVTGVCETLYEIHPYPEYVLQ----SKPWVVPHKH 46
XP_044257336 TQ-ED-SNTAVFKTMEETVTGVTETLYEIHPYPEYVLQ----SKPWVVPHKH 46
CAH0562014   PE-GQ-QNNAVFKTMEDTINGQSETLYDIHALPEYVLQ----SKPWLAPKHE 46
KAH1029837   PN-DE-KMEAVFQTMEDTVTGETETNYDIHQLPEYLVD----SQPWLVPQQQ 46
CAH0562016   PE-GQ-QNNAVFKTMEDTINGQSETLYDIHALPEYVLQ----SKPWLAPKHE 46
ERL95541     PN-DE-KMEAVFQTMEDTVTGETETNYDIHQLPEYLVD----SQPWLVPQQQ 46
CAH0562021   PE-GQ-QNNAVFKTMEDTINGQSETLYDIHALPEYVLQ----SKPWLAPKHE 46
CAG9759825   PS-HG-RMDGVFKTMEETITGESETLYDIHPLPEYLLQ----SKPWLAPQQD 46
KAJ8923586   PEDDD-QNSAVFKVMEETVTGKIETTYDIHPLPEYILQ----SEPWLVRYPD 47
XP_048525023 PN-DE-KMEAVFQTMEDTVTGETETNYDIHQLPEYLVD----SQPWLVPQQQ 46
QQM99833     PK-EG-SNNAVFKTLEDTITGITETTYDIHPLPEYILQ----SEPHLAPFPH 46
KAJ8936451   PE-DQ-GNSAVFKTMEETITGNTETLYDIHPIPEYVLQ----SKPWMARHLE 46
XP_018565741 PE-DD-QNSAVFKTMEDTVTGKTETMYDIHPMPEYVLQ----SKPWLARHLE 46
RZC42857     SQ-EG-SNTAVFKTMEETVTGISETLYEIHPFPEYVLQ----TYPWLVPLDN 46
XP_044257065 TQ-EG-SNTAVFKTMESTVTGESETLYEIHPLPEYILQ----TTPWLVPLKH 46
XP_970210    TQ-EG-SSTAVFKTMEPTVTGESETLYEIHPLPDYILQ----TTPWLAPLKH 46
XP_019873436 PD-MT-TNTANFKTMEDTVTGNTETLYDIHLLPEYVLQ----NQPWLAPKYE 46
XP_030756598 PS-GD-NLSGVFKTMEKTVTGETETEYKIHNLPLYILQ----SRPWLAPKSD 46
XP_060515924 PS-EG-RNDGVFKTLEETITGETVTSYEIHPLPEYQLQ----SEPWLAPQLN 46
KAJ8985867   ---GN-EERYVFNVMEETVTGNTETMYDIHPIPEYTFQ----NAPLMSRLLK 44
KAH0815682   TQ-DD-SNTAVFKTTEETVTGVAETLYEIHPYPNYVLQ----TTPWVVPQKE 46
CAG9828227   PEEGPNNQEAVFLAMEDTVTGETETLYLIRPLPEEVVQ----RKSGEAKYLD 48
AMK38869     SE-NQ-QESAVFRIMEDTVTGNTDTLYEIKQLPEYMLQ----SKPWEARHLQ 46
QGN03672     SQ-DY-SNSAVYKTMEETVTGTCETLYEIHPYPQHVLP----SKPYVAPHKR 46
CAH1377764   TQ-EG-SNNAVFKTMEKTVTGEAETLYEIQPLPDYILQ----TTPWLAPLRD 46
CAG9759824   PK-NN-EMEGVFKSMEETVTGETETLYQIHRLPEYLTQ----SQPWLVPQQQ 46
UXX33469     PE-KG-ESNGIFKTMEETVTGKYETLYELNPLPKYILQ----SEPQWVPLPE 46
UVT84831     PE-EG-SNYGVYKTMEDTVTGRYETLYEINPIEGYMAR---------GLKEE 41
XP_045482383 PN-DE-TNNAVYKTIEDTVTGITETSYDIHPMPEYILQ----AEPHLAPFPH 46
XP_050313480 PT-NE-YMDAVFKTMEETVTGKTETIYDIHRLPEYLVQ----SQPWIAPQYK 46
KAG5892825   LQ----HESAVFTVMEDTVTGNSDTLYEVRPLPKHLLQ----SKPWEIRLVD 44
XP_023016640 PD-GE-KNSAIFVTMEDTVSGKTSTLYEIRPIPKHVVQ----YNKQESKYLD 46
Q05808       PT-NE-YMDAVFKTMEETVTGKTETIYDIHRLPEYLVQ----SQPWIAPQYK 46
XP_063928892 PQ-DD-SETAVFKTMEKTVTGESETLYEIHPFPEYLLQ----TEPSLAPKKH 46
XP_044765433 PK-HG-SNNAVFKTVEDTVTGVTETSYDIHPMPEYILQ----SEPHLAPFPH 46
APR62727     PK-EG-SNNAVYKTVEDTITGITETTYDIHPMPEYMIQ----SKPYLAPYPH 46
KAI4462317   PE-GD-DIDAVFKTMEETVTGRVETMYELNPLPEHILQ----SHPELVPMDH 46
XP_044765434 PK-EG-SNNAVFKTLEDTVTGITETNYDIHPMPEYMLQ----SKPYLAPYPH 46
KAI4462319   PE-GD-DIDAVFKTMEETVTGRVETMYELNPLPEHILQ----SHPELVPMDH 46
KAJ3622909   TQ-ED-SNTAVFKTTEETVTGVAETLYEIHPYPNYVLQ----TTPWVVPQKE 46
KAG5880947   DE----PSYDVFVCMEDTISGNTETLYEIRPLPDYELD----NREYESRYLD 44
CAH1377765   TQ-DD-SNTAVFKTTEETVTGVAETLYEIHPYPNYVLQ----TTPWVVPQKE 46
AAU20328     TQ-DD-SNTAVFKTTEETVTGVAETLYEIHPYPNYVLQ----TTPWVVPQKE 46
KAG5880948   RP-DE-PIYDVFVCMEDTISGNTETIYEIRPLPEYVLG----NREYESRYLD 46
XP_019765359 PE----RFDAVFKTMEETVTGKTETIYEIHPLPDYLIQ----SQPWLAPQQK 44
CAH1153579   PT-AD-GNSAVFATMEDTVTGNTRTLYEIRPLAPHVLQ----DKYSTLRFAQ 46
ALN38803     PS-ED-KLDGVFKTMEETVTGETETTYKMHPLPLYILQ----SQPWLVRKDD 46
XP_028134501 PEDGPNKQEAVFVAMEDTLTGKTETLYLIRPLPDQVVQ----RKSSEAKYLD 48
VEN45879     ET----GIPSVFKTMEETVTGNTETLYEINELPEYLLQ----SEEYDYLKDR 44
XP_018322689 PK-EG-SDFAVFKTMEKTITGNTETLYEIKQYPEYLLQ----WENWRESLPH 46
AMK38870     PS-GE-HNSAVFTTMEETVSGNTQTLYEIRPLPDYVLQ----NREHESRYLD 46
AKR04341     PE----EHATHFQVVEDTVTGETETIYKIFRLPEHMIQ----QEPQAERLMQ 44
CAG9828226   PE-HG-SNSGVFLTMEDTILGETETLYKIRPVPEHTLLTSEQEARFVELKQK 50
CAG9816778   SQ----EQPAVFKVLEDTVTGNTETLYEIREVPEYLIQ---NDEPWQQRLFR 45
VEN45878     ET----GIPSVFKTMEETVTGNTETLYEINELPEYLLQ----SEEYDYLKDR 44
CAH2002692   EG----GLPSVFKTMEETVTGNTETMYEINELPEYLLQ----DEEFDYLKDR 44
QDP42284     PHNEDDDNDAVFFTKEDTVTGNVETLYQIRPLPDHIVQ----GNPDEANYLD 48
CAH1103550   PENGDFDNDAVFMTMEDTITGNTETMYQIRPLSDHLVQ----ADFKEAQYLD 48
XP_023017897 SK-NR-FEPTVFSVMEDTITGNSDTLYEVKIKPRFMFE----SRPWEIRMLQ 46
XP_028134500 PE-HG-SNSGVFLTMEDTILGETETLYKIRPVPEHTLLTNEKEARFVELKQK 50
XP_028134505 ---DDRPDSAMFRIMEETVSGNTETLYEINPMPEHLMV-----NSDLVPDFQ 44
CAG9855083   PENGDGDNDAVYTAMEDTVTGNTATLYQIRPLPAHLVQ----ADPEEAEYVD 48
KAK4872055   PE-NN-TFAGVFKTMEDTVTGKYATLYELNELPSVATQ----ARPWLVPYPQ 46
CAH1103551   PT-SQ-SNNAVFLTMEDTVSGETETLYEIRPVSEYQFD-----FAKEGKHFQ 45
WFV29669     PE-NG-DNDAVYVVMEDTVTGETETIYNIRPLPDYIVQ----SRNNEEKYLD 46
CAG9828225   ---DDRSDSAMFRLMEETVSGNTETLYEINPIPAHLLV-----NSGIVPDFQ 44
CAG9855084   PS-PE-SNDAIFMAMEDTVSGETETLYEIRPVPQHLFD----ETEKHKNFQQ 46
XP_031335608 PE-ND-SYSGVYKTMEDWVTGKFATLYEMNEVPSTQIQ----YKPWRVPFPQ 46
WLD26188     PD-PE-SNSGVFLTKEDTVLGVTETLYKISPIPEHLLFNSEENARLVDFKLK 50
WAP81825     PK-EG-SNNAVFKTLEDTVTGITETNYDIHPMPEYMLQ----SKPYLAPYPH 46
XP_057657771 PEDGDWDNDAVYLVMEDTVSGEVETTYNIRPLPDHVVQ----SRSNEARYLD 48
CAH1103552   TQNDNREDSAFYKVYEETVSGNTETVYEINPIPEYKLF------DSEVPQLQ 46
XP_056645845 PEDGDWDNDAVYLVMEDTVSGEVETTYNIRPLPDHVVQ----SRSNEARYLD 48
QDP42283     PN-DN-SNNAIFMAMEDTVSGQTETLYEIRPVPEHLIQGKLDQEGLEFPQ-- 48
CAG9855085   ---DNSQNSAYFKTFEETVTGNTETLYEVSPFPQYALN----SLNYEIPRLQ 45
KAF5287239   PK-QN-TFPAVFKTMESTTVGKYSTLYEVKVLPDVVTQ----DRPELIPYPQ 46
AMP18919     PN-DE-TNNAVYKTIEDTVTGITETSYDIHPMPEYILQ----AEPHLAPFPH 46
WLD26187     TEKGDRHDSASFIIPEETVSGHTDTFYEINPLPEHLLL-----NSAIVPDFQ 47
QDP42282     ---KGNQNSAKFQVYEETVTGHTDTMYEVSPLPEFKTF-----DESIPEFQR 44
CAH2002694   EG----GLPSVFKTMEETVTGNTETMYEINELPEYLLQ----DEEFDYLKDR 44
CAH2002693   EG----GLPSVFKTMEETVTGNTETMYEINELPEYLLQ----DEEFDYLKDR 44
XP_056646762 ------ADPDSFIIPEETVSGTTDTSYKINRIPEYLLL-----NSEIVPDFQ 41
KAF5294837   PK-SG-SSFAIFKTAEETITGKYPTVYELNELTPAEIQ----RRAWKVPYPQ 46
CAH1153577   ---DE-YNLMNFKVMEETITGKVETHYEVKQIPKKYVS----T-------FD 37
CAH2002695   EG----GLPSVFKTMEETVTGNTETMYEINELPEYLLQ----DEEFDYLKDR 44
```

*Hemiptera* (Vg)

```
                                              β-sheet
                                    predicted peptides
AAB72001       NLKDSKINLV-PSGQSG-EPMGVYKTMEDSVNGIYETLYDISVLPEYILQA 49
AFW97644       NLKSSRINSL-PNKD---QAYGVYKTMEDTVAGEVETIYDISPLPQYVIQS 47
AGJ26477       NAIKSRRNIL-PQSDSNQQVSGSFKAMEDSVTGKCETHYDVDELPMRVVQQ 50
AGJ26478       NAIKSRRNIL-PQSDSNQQVSGSFKAMEDSVTGKRETHYDVDELPMRVVQQ 50
AGT39945       NLKSSRINSL-PSKD---QAYGVYKTMEDTVAGEVETIYDISPLPQYVIQS 47
AGV05363       NLQKSRINSL-PSKD---QAYGVYKTMEDTVAGEVETIYDIPPLPQYIIQS 47
AIA09041       NLKKSRVNQV-PVEG---QTIGVYKTVEDTVTGECETIYDISPLPQYVLQS 47
AJI43739       NLKSSRINSL-PNGN---QAYGVYKTMEDTVAGEVETIYDVSPLPTYVIQS 47
ALN70475       NLKKSKINLL-PKDQHK-GKMAVFKVMEDSVNGIYETTYDISPMPEYVIQS 49
AOY34570       NLQKSRINSL-PDRE---NVNGVYKTMEDSVSGECETLYDISPLPKVVLQN 47
AQM52239       NEVSSRLNQK-PKNG---KNFGTFKTMEDTVTGETETLYDIKPLEQVEYQN 47
BAA85987       NIQKSRSNSLFDNSD---DPVVTFKTMEASVTGKCETLYDFSPLLKQELME 48
BAA88075       NLKKSKFNDV-PKGN---QLTGIFKTWEDSVNGEYEVMNEVSVLPEYLIES 47
BAA88076       NLKESKLNSV-PHGE---QDSGVFTTMEESTNGKYETIYEVSVLPEYILQS 47
BAA88077       NLKQSKLNNV-PQGG---QQNGVYKTYEDSINGVYETIYEFYPLPEYVLQS 47
BAG12118       NLQKSRINQL-PAMH---KPMGVYKTMEDTVTGECETVYDVSPLPDYLLQS 47
BAJ33507       NLKNSRINSL-PNKN---QAYGVYKTMEGTVAGEVETIYDISPLPQYVIQS 47
BAP87098       NAIKSRRNIV-PNGQ---QVSGSFKVMEDSVTGKCETHYDVDELPMRVVQQ 47
BAU36889       HLMKSKINQV-PNKG---DHMGVYKTMEDTVTGLCETIYDVSPLPEYVLQS 47
BAU68162       NLKKSKINDV-PKDN---QHTGIYKTMEDSINGEYETIYDISVLPEYLLES 47
BES91476       NLQKSRINSL-PSKD---QAYGVYKTMEDTVAGEVETIYDISPLPQYIIQS 47
BES91478       NLKNSRINSL-PSKD---QAYGVYKTMEDTVAGEVETIYDISPLPQYIIQS 47
BES91480       NLKNSRINSL-PSKD---QAYGVYKTMEDTVAGEVETIYDISPLPQYIIQS 47
CAH1388802     NVIKSRLNLI-PNSD---LSLGTYKTMEDSVNGVHETIYDISEIQQYALQS 47
CAH1388803     NVIKSRLNLI-PNSD---LSLGTYKTMEDSVNGVHETIYDISEIQQYALQS 47
CAH1388807     NLKYSKLNNV-PEDN---QDSAIFSTMEECINGKYETIYDVSVIPEYILQS 47
CAH1388808     NLKHSKVNSL-PQGD---QQSRAYKTKEGSVSGVDETLHEIYRMPEYIIQS 47
KAF6215813     NLKKSSYNQI-PQSN---DPTASFKTTEDTVHGEVETSYSLSPLPTYHLHS 47
KAF6216356     NLKSSRINSL-PNKD---QAYGVYKTMEDTVAGEVETIYDISPS------- 40
KAG8258925     NLQKSRINSL-PTQG---SVNGVYKTMEDSVSGECETLYDISPLPKLVLQN 47
KAG8258927     NLQKSRINSL-PTQE---TVNGVYKTMEDSVSGECETLYDISPLPKVVLQN 47
KAI5699898     NEVSSRLNQK-PKNG---KPFGTFKTMEDTVTGECETLYDIKPLQQVEQQN 47
KAI5700205     NEVSSRLNQK-PKNG---KPFGTFKTMEDTVTGECETLYDIKPLQQVEQQN 47
QDD67294       NVIKSRRNIV-PNSN---QVSGSFKAMEDSVTGKCETHYDVDLLPMRVVQE 47
QDQ29854       NLKKSKINQV-PSKG---QAMGVYKTMEDTVTGECETIYDISPLPQYVLQS 47
QFQ33312       KINKKDKSNYSSKSKSRPDLEDSFTVMQETVVGKCEVQYEVFRLPRNGEGW 51
QHB15616       NLKHSSHNQL-PKVN---KPYGVYKTMEDSVTGECETLYDVSPLPEITLQT 47
QHD25542       NVIKSRRNIV-PNSN---QVSGSFKAMEDSVTGKCETHYDVDLLPMRVVQE 47
QIQ19557       KNNKDKSSLS-SKSESRLDVENSFTVMEETVVGKCEVQYEVFITPRNGEGW 50
QIQ19558       NVIKSRRNIV-PNSN---QVSGSFKAMEDSVTGKCETHYDVDLLPMRVVQE 47
QXD38625       HLKPSHFNDV-PKGE---QQTGAFKTMEDCVSGRYEVLYDVGVLPPYILQS 47
QXD38626       NLKKSIWNNV-PNGE---QQNGVFTTKEGSVHGISETLYQINPMPEYLVQS 47
QXD38627       NLKKSKWNNV-PSGK---QMSGIYKTKEDSVNGVYETCYEVYPMQEYVLQS 47
RZF36490       NAIKSRRNIL-PQSDSNQQVSGSFKAMEDSVTGKCETHYDVDELPMRVVQQ 50
RZF38098       KSNKDKSSLS-SKSDSRQDLEDSFTVMEETVVGKCEVQYEVFISPRNGEGW 50
UOL49140       NEVSSRLNQK-PKNG---ENFGTFKTMEDTVTGETETLYDIKPLEQVEYQN 47
WLK26396       NIKDSRFNLV-PRSE---MGTGAYKIMEDSIHGIYETIYDVSLLPEILLQS 47
XP_014270480   NVIQSQMNQI-QNSD---ISISNYKTMEDSVNGVYETIYDISEIPEYILQS 47
XP_014270535   NLKESKFNSV-PHGE---QDSGTFSTMEESIHGKYETNYVVSVLPEYLLQS 47
XP_014270538   NLQHSKLNNV-PQGK---QQNGVYKTKEDSINGIYETTYDVNPMPYYILQS 47
XP_014291484   NLKKSKINDV-PKGQ---QQTGIYKTMEDSINGEYETIYEISDIPAYIVQS 47
XP_014293026   NLKKSKINDV-PKGQ---QQTGIYKTMEDSINGEYETMYEISVMPEYLLSS 47
XP_026682480   NEVSSRLNQK-PKNG---KPFGTFKTMEDTVTGECETLYDIKPL------- 40
XP_026682481   NEVSSRLNQK-PKNG---KPFGTFKTMEDTVTGECETLYDIKPLQQVEQQN 47
XP_039298390   NAIKSRRNIV-PNGQ---QVSGSFKVMEDSVTGKCETHYDVDELPMRVVQQ 47
XP_039298391   NAIKSRRNIV-PNGQ---QVSGSFKVMEDSVTGKCETHYDVDELPMRVVQQ 47
XP_039298807   KINKKDKSNYSSKSKSRPDLEDSFTVMQETVVGKCEVQYEVFRSPRNGEGW 51
XP_046672836   NLQKSRINSL-PTQG---SVNGVYKTMEDSVSGECETLYDISPLPKLVLQN 47
XP_046674730   NLQKSRINSL-PTQE---TVNGVYKTMEDSVSGECETLYDISPLPKVVLQN 47
XP_054267470   NLQKSRINSL-PNKE---NVNGVYKTMEDSVGGECETLYDISPLPKVVLEN 47
XP_054269933   NLQKSRINSL-PNKE---NVNGVYKTMEDSVGGECETLYDVSPLPKVVLEN 47
XP_054280083   SRPGHYLNQL-TNT----QTSDVYSVFEDSVSGRCEVLYEINPLPKQQGPD 46
```

*Thysanoptera* (Vg)

β-sheet

predicted peptides

```
XP_026289649.2 AENIISSRINNKPSHGSVNGVYKTMEDDVTGECEVVYDISPLPQYQLQSRPELAPMPQ 58
KAE8740093.1   AENLISSRINNKPSHGSLNGVYKTMEDSVTGECEVMYDISPLPQYQLQSRPELAPMPQ 58
XP_052120755.1 AENLISSRINNKPSHGSVNGVFKTMEDDVTGECEVMYDISPLPQYQLQSRPELAPMPQ 58
XP_034239803.1 AENLISSRINNKPSHGSANGVYKTMEDSVTGECEVMYDISPLPQYQLQSRPELAPMPQ 58
KAK3914139.1   AENLISSRINNKPNHGSVNGVYKTMEDSVTGECEVMYDISPLPQYQLQSRPELAPMPQ 58
KAJ1522457.1   AENLISSRINNKPSHGSLNGVYKTMEDSVTGECEVMYDISPLPQYQLQSRPELAPMPQ 58
KAJ1522462.1   AENLISSRINNKPSHGSLNGVYKTMEDSVTGECEVMYDISPLPQYQLQSRPELAPMPQ 58
KAK3925741.1   AENLISSRINNKPSHGSVNGVFKTMEDDVNGECEVMYDISPLPQYQLQSRPELAPMPQ 58
KAK3926221.1   AENLISSRINNKPSHGSVNGVYKTMEDSVTGECEVMYDISPLPQYQLQSRPELAPMPQ 58
XP_034253661.1 AENIISSRINNKPSHNSVNGVFKTMEDDVTGECEVMYDISPLPQYQLQSRPELAPMPQ 58
KAK3912473.1   AENLISSRINNKPSHGSVNGVFKTMEDDVNGECEVMYDISPLPQYQLQSRPELAPMPQ 58
XP_026272532.2 AENLISSRINNKPSHGSLNGVYKTMEDSVTGECEVMYDISPLPQYQLQSRPELAPMPQ 58
XP_034256232.1 AENLISSRINNKPSHNSVNGVFKTMEDDITGECEVMYDISPLPQYQLQSRPELAPMPQ 58
KAK3917678.1   AENIISSRINNKPSHNSVNGVYKTMEDDVTGECEVIYDISPLPQYQLQSRPELAPMPQ 58
KAE8749850.1   AENLISSRINNKPSHGSVNGVFKTMEDDVTGECEVMYDISPLPQYQLQSRPELAPMPQ 58
```

### *Hymenoptera* (Vg)

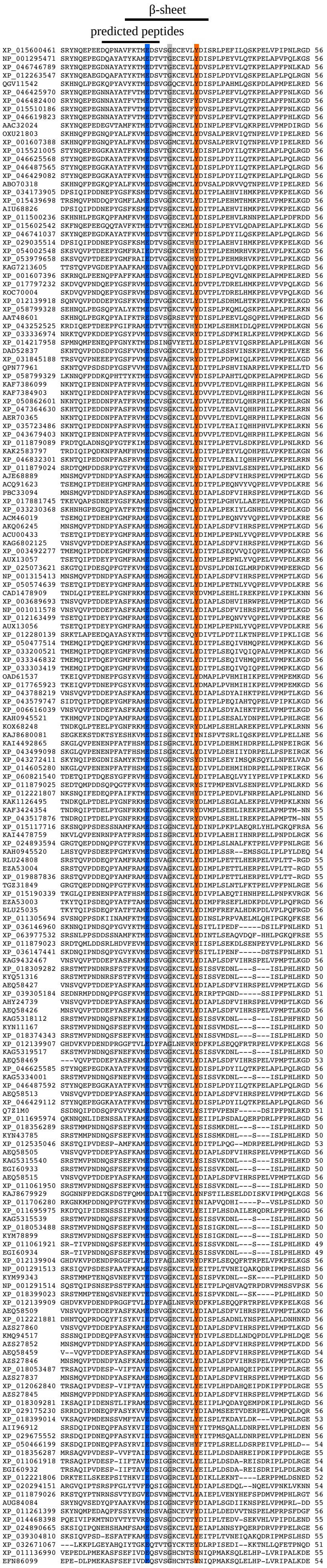

*Lepidoptera* (Vg)

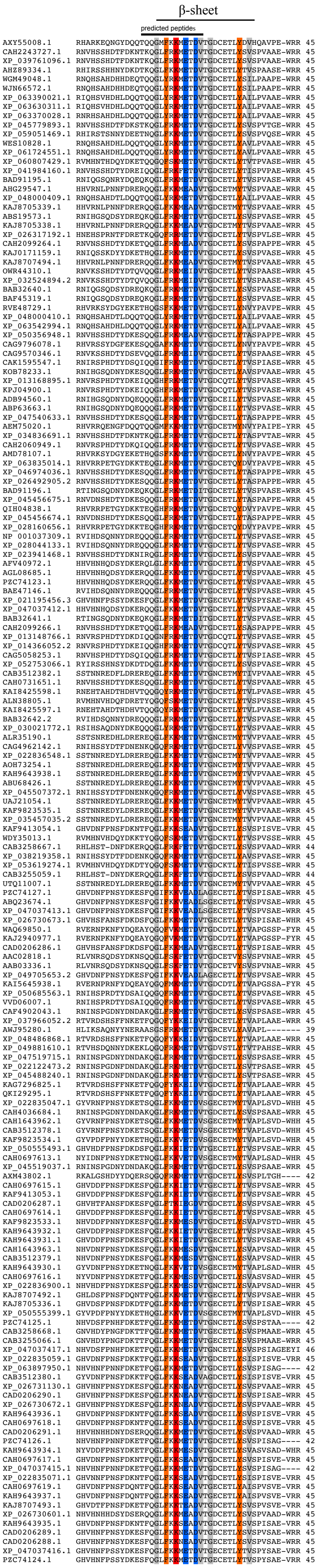

### *Diptera- non Brachycera* (Vg)

β-sheet

predicted peptides

```
KAG4067618.1   NEIKCKHNQLPEK-----DGNNAVYKTMEPTVTGKCETIYDISPLPEY-LTQSHRDLVPMPQLKGTG 61
XP_037043261.1 NEIKCKHNQLPEK-----DGNNAVYKTMEPTVTGKCETIYDISPLPEY-LTQSHRDWVPMPQLKGTG 61
KAJ6633698.1   NEIKCKYNQLPEK-----DSNNAVFKTMEATVTGKCETIYDISPLPEY-LTQAHREWVPMPQLKGQG 61
XP_037038024.1 NEIRCKHNQLPEE-----HGNNAVYKTMEPTVTGECETLYDISPLPEY-LTQNHRDWVPMPQLKGNG 61
CRK93348.1     YLIKCSSNQLPEP-----KTVNGVFKTMEPSVTGKCETMYDVSPIPEH-KVASNPSWVPLPQLKREH 61
KAG5666818.1   NLIKCRRNQMPDY-----ERSSGVYKTMEPSVTGKCQTMYDISPLPAY-QVASNPHWAPMPHLEHED 61
CAG9810144.1   NLIKCRRNQLPNH-----ETHTAAYKTMEPSVHGKCKTMYDISPIPDY-QVAANQEWAPMPSLKKDG 61
XP_063702688.1 NEIKCDKNEFPEK-----EDMTGVFKVMERAVTGECETLYDVSPLPRY-MLQSHPHWAPMPELNKEK 61
ADH04225.1     NVVKSKYNQFPEN-----TTFTGVYKVMEPLVTGECETLYDVNVVPEH-VIKGNKYFVPRPDLREEN 61
XP_039447468.1 NVIKSKYNQFPEN-----TTFTGVYKVMEPLVTGECETLYDVNVVPEH-VIKGNKRFVPRPELREDN 61
XP_001857970.2 NVIKSKYNQFPEN-----TTFTGVYKVMEPLVTGECETLYDVNVVPEH-VIKGNKRFVPRPELREDN 61
AAV31930.1     NVIKSKYNQFPEN-----TTFTGVYKVMEPLVTGECETLYDVNVVPEH-VIKGNKRFVPRPELREDN 61
XP_055383870.1 NLIKSRHNQLPQD----KHNPTALYKTMEPTVTGECETIYDISPLQEAIVEQTEAQWAPKVHLNAEG 63
XP_063701775.1 NLIKCRHNQLPQK-----GLYNAYYKVMEPSVTGNCETAYTVSPFPKY-LIQSHPQLAPFPSLAHND 61
XP_058450291.1 NKINSKYNQLPEN-----NSFTGAYKVMEPSVTGNCETLYDVNAVPEY-IAQSNPEWVPQPQLREDG 61
XP_055607086.1 NLIKSKYNQLPEN-----ETITAVYSTMEPSIAGECATLYDINAIPKY-KIQSHKEWVPQPQYLQEN 61
XP_058450293.1 NKINSKYNQFPEN-----NSFTGAYKVMEPSVTGNCETLYDVNAVPEY-MTRSYPEWVPQPQLREDG 61
XP_055384020.1 NLIKSRHNQLPQD----KHNPTALYKTMEPTVTGECETIYDISPLQEAIVEQTEAQWAPKVHLNAEG 63
XP_058824326.1 NKMNSKYNQFPEN-----NSFTGVYKVMEPTVTGNCETLYDVNAVPEY-IAQANPDWAPKPQYREEG 61
XP_058824918.1 HAINSKYNQQPEN-----NSLTGAYKVMEPVVTGNCETMYDVNVVPEY-ISQANKEWVPKPELREDG 61
XP_055541618.1 NEIDYKYNQLPEN-----NSFTGVYKVMEPVVTGNCEMLYDVNAVPEY-LMQSYKKFAPQPHFREEG 61
XP_055609491.1 NLIKSKYNQLPEN-----NTLNGVYKVMEPVVTGECETLYDINPIQPF-MVKANRQWVPQPQLREDG 61
AAF82132.1     YVIKSEFNQFPEN-----NTLTGVYKTMEPSVTGECETLYDVNPVPEF-HFQSHKEWVPQPQWLEED 61
XP_052870874.1 YVIKSEFNQFPEN-----NTLTGVFRTMEPSVTGECETLYDVNAVPEY-FIQSHNEWVPQPEYFEDY 61
XP_055609492.1 NLIKSKYNQLPEN-----NTLNGVYKVMEPVVTGECETLYDINPIQPF-MVKANRQWVPQPQLREDG 61
XP_039429667.1 NLIKSKYNQLPEN-----ETANAVYKTMEPSVSGECETLYDVNVLPKY-KIQSHDEWVPQPQYMQQD 61
XP_039429669.1 NLIKSKYNQLPEN-----ETANAVYKTMEPSVSGECETLYDVNVLPKY-KIQSHDEWVPQPQYMQQD 61
EDS30760.1     NLIKSKYNQLPEN-----ETANAVYKTMEPSVSGECETLYDVNVLPKY-KIQSHDEWVPQPQYMQQD 61
XP_058064674.1 YVIKSEFNQFPEN-----NTLTGVFKTMEPSVTGECETLYDVNAVPEY-FIQSHKEWVPQPEYFEED 61
EDS30759.1     NLIKSKYNQLPEN-----ETANAVYKTMEPSVSGECETLYDVNVLPKY-KIQSHDEWVPQPQYMQQD 61
XP_038117787.1 NLIKSKYNQLPEN-----ETANAVYKTMEPSVSGECETLYDVNVLPKY-KIQSHDEWVPQPQYMQQD 61
ADH04227.1     NLIKSKYNQLPEN-----ETANAVFKTMEPSVSGECETLYDVNVLPKY-KIQSHDEWVPRPQYMQQD 61
XP_058064675.1 YVIKSEFNQFPEN-----NTLTGVFKTMEPSVTGECETLYDVNAVPEY-FIQSHKEWVPQPEYFEED 61
XP_055609493.1 NLIKSKYNQLPEN-----NTLNAVYKVMEPVVTGECETLYDINPIQPF-MIKANRQWVPQPQLREDG 61
XP_052870875.1 YVIKSEFNQFPEN-----NTLTGVFRTMEPSVTGECETLYDVNAVPEY-FIQSHNEWVPQPEYFEEY 61
AAV31931.1     NLIKSKYNQLPEN-----ETANAVYKTMEPSVSGECETLYDVNVLPKY-KIQSHDEWVPQPQYMQQD 61
EGK97015.1     YVIKSEFNQFPEN-----NTLTGVYKTMEPSVTGECETLYDVNPVPEF-HFQSHKEWVPQPQWLEED 61
XP_001843135.2 NLIKSKYNQLPEN-----ETANAVYKTMEPSVSGECETLYDVNVLPKY-KIQSHDEWVPQPQYMQQD 61
EDS34782.1     NVIKSKYNQFPEN-----TTFTGVYKVMEPLVTGECETLYDVNVVPEH-VIKGNKRFVPRPELREDN 61
XP_001857967.2 NVIKSKYNQFPEN-----TTFTGVYKVMEPLVTGECETLYDVNVVPEH-VIKETSASSRDP-LREDN 60
XP_035895626.1 YVIKSEFNQFPEN-----NTLTGVYKTMEPSVTGECETLYDVNPVPEF-HFQSHKEWVPQPQWLEED 61
ABD97990.1     YVIKSEFNQFPEN-----NTLTGVYKTMEPSVTGECETLYDVNPVPEF-HFQSHKEWVPQPQWLEED 61
ADH04226.1     NLIKSKYNQLPEN-----ETANAVFKTMEPSVSGECETLYDVNVLPKY-KIQSHDEWVPRPQYMQQD 61
XP_053680606.1 YVIKSEFNQFPEN-----NTLTGVFKTMEPSVTGECETLYDVNAVPEF-HFQSHKEWVPQPQWLEED 61
XP_035895623.1 YVIKSEFNQFPEN-----NTLTGVYKTMEPSVTGECETLYDVNPVPEF-HFQSHKEWIPQPQWLEED 61
XP_053680330.1 YVIKSEFNQFPEN-----NTLTGVFKTMEPSVTGECETLYDVNAVPEF-HFQSHKEWVPQPQWLEED 61
XP_041767428.1 YVIKSEFNQFPEN-----NTLTGVYKTMEPSVTGECETLYDVNPVPEF-HFQSHKEWVPQPQWLEED 61
AGZ02020.1     YVIKSEFNQFPEN-----NTLTGVYKTMEPSVTGECETLYDVNPVPEY-HFQSHKEWVPQPQWLEED 61
XP_052903498.1 YVVKSEFNQFPEN-----NTLTGVYKTMEPSVTGECETLYDVNPVPEF-HFQSHKEWVPQPQWLEED 61
XP_058130499.1 YVIKSEFNQFPEN-----NTLTGVFKTMEPSVTGECETLYDVNAIPEY-FLQSHKEWVPQPQYLEED 61
XP_058130501.1 YVIKSEFNQFPEN-----NTLTGVFKTMEPSVTGECETLYDVNAIPEY-FLQSHKEWVPQPQYLEED 61
XP_052903497.1 YVVKSEFNQFPEN-----NTLTGVYKTMEPSVTGECETLYDVNPVPEF-HFQSHKEWVPQPQWLEED 61
XP_050067904.1 YVIKSEFNQFPEN-----NTLTGVYKTMEPSVTGECETLYDVNPVPEY-HFQSHKEWVPQPQWLEED 61
XP_050067906.1 YVIKSEFNQFPEN-----NTLTGVYKTMEPSVTGECETLYDVNPVPEY-HFQSHKEWVPQPQWLEED 61
XP_050083776.1 YVIKSEFNQFPEN-----NTLTGVFKTMEPSVTGECETLYDVNAVPEY-FIQSHKEWVPQPQYFEED 61
XP_061498654.1 YVIKSEFNQFPEN-----NTLTGVYKTMEPSVTGECETLYDVNPVPEF-HFQSHKEWVPQPQWLEED 61
XP_035774741.1 YVIKSEFNQFPEN-----NTLTGVFKTMEPSVTGECETLYDVNAVPEY-FIQSHKEWVPQPQYYEED 61
XP_049462553.1 YVIKSEFNQFPEN-----NTLTGVYKTMEPSVTGECETLYDVNPVPEF-HFQSHKEWVPQPQWLEED 61
XP_050083777.1 YVIKSEFNQFPEN-----NTLTGVFKTMEPSVTGECETLYDVNAVPEY-FIQSHKEWVPQPQYFEED 61
XP_061498653.1 YVIKSEFNQFPEN-----NTLTGVYKTMEPSVTGECETLYDVNPVPEF-HFQSHKEWVPQPQWLEED 61
XP_061498652.1 YVIKSEFNQFPEN-----NTLTGVYKTMEPSVTGECETLYDVNPVPEF-HFQSHKEWVPQPQWLEED 61
XP_040160637.1 YVIKSEFNQFPEN-----NTLTGVYKTMEPSVTGECETLYDVNPVPEF-HFQSHKEWVPQPQWLEED 61
XP_040220166.2 YVIKSEFNQFPEN-----NTLTGVYKTMEPSVTGECETLYDVNPVPEF-HFQSHKEWVPQPQWLEED 61
AAF82131.1     YVIKSEFNQFPEN-----NTLTGVYKTMEPSVTGECETLYDVNPVPEF-HFQSHKEWVPQPQWLEED 61
XP_040160603.1 YVIKSEFNQFPEN-----NTLTGVYKTMEPSVTGECETLYDVNPVPEF-HFQSHKEWVPQPQWLEED 61
XP_040220168.2 YVIKSEFNQFPEN-----NTLTGVYKTMEPSVTGECETLYDVNPVPEF-HFQSHKEWVPQPQWLEED 61
EAA08615.3     YVIKSEFNQFPEN-----NTLTGVYKTMEPSVTGECETLYDVNPVPEF-HFQSHKEWVPQPQWLEED 61
XP_049462552.1 YVIKSEFNQFPEN-----NTLTGVYKTMEPSVTGECETLYDVNPVPEF-HFQSHKEWVPQPQWLEED 61
XP_053660489.1 YVVKSEFNQFPEN-----NTLTGVYKTMEPSVTGECETLYDVNPVPEF-HFQSHKEWVPQPQWLEED 61
UOW66183.1     YVIKSEFNQFPEN-----NTLTGVYKTMEPSVTGECETLYDVNAVPEF-HFQSHKEWVPQPQWLEDD 61
AHN13887.1     YVIKSEFNQFPEN-----NTLTGVYKCMEPSVTGECETLYDVNPVPEY-HFQSHKEWVPQPQWLEED 61
AHM10344.1     YVIKSEFNQFPEN-----NTLTGVYKTMEPSVTGECETLYDVNPIPEF-HFQSHKEWVPQPQWLEED 61
XP_035774740.1 YVIKSEFNQFPEN-----NTLTGVFKTMEPSVTGECETLYDVNAVPEY-FIQSHKEWXPQPQYYEED 61
XP_049282984.1 YVIKSEFNQFPEN-----NTLTGVYKTMEPSVTGECETLYDVNPVPEY-HFQSHKEWVPQPQWLEEN 61
XP_049533828.1 YVIKSEFNQFPEN-----NTLTGVFKTMEPSVTGECETLYDVNAVPEY-FIQSHKEWVPQPQYFEED 61
ETN67132.1     YVIKSEFNQFPEN-----NTLTGVFKTMEPSVTGECETLYDVNAVPEY-FIQSHKEWVPQPQYFEED 61
AAV31933.1     YVIKSEFNQFPEN-----NTLTGVFKTMEPSVTGECETLYDVNAVPEY-FIQSHKEWVPQPQYYEED 61
XP_049282983.1 YVIKSEFNQFPEN-----NTLTGVYKTMEPSVTGECETLYDVNPVPEY-HFQSHKEWVPQPQWLEEN 61
XP_049533827.1 YVIKSEFNQFPEN-----NTLTGVFKTMEPSLTGECETLYDVNAVPEY-FIQSHKEWVPQPQYFEED 61
AEO51020.1     YVIKSEFNQFPEN-----NTLTGVYKTMEPSVTGECETLYDVNPVPEY-HFQSHKEWVPQPQWLEED 61
XP_062555811.1 NMYKSKHNQFPGN-----NSFTGMYEVMEPVVTGKCKTMYDVSVLPEY-MIKANSHMVPQPQLREDD 61
AAV31932.1     NAIKSKYNQFPEN-----NSFTGVYKVMEPQVTGHCETLYDVKVVPEY-LVQANKQWVPQPQFREKN 61
XP_055922367.1 NVIKCRHNQLPRE---SAGHTNALYKTMEPTVTGECETLYEMTTIPAFSVTEQTEKWARDTQLNE-- 62
XP_055637489.1 NAIKSKYNQFPEN-----NTFTGVYKVMEPQVTGHCETMYDVKVVPEY-LVQANKQWVPQPQFREEG 61
AAV31926.1     HMYKSKYNQVPGN-----NSFTGMYEVMEPVVTGKCKTLYDVSVIPPY-MVQANKHWVPLPQLREEG 61
XP_055312945.1 NAMESQDNHLPQNQDNEDNHNQALFKVMEPTVTGKCESFYDISRVPLY-LAQAYPEYDSNVQLQQGE 66
XP_055922280.1 NVIKCRHNQLPGE---SAGHTNALYKTMEPTVTGECETLYEMTTIPAFSVTEQTEKWARDTQLNE-- 62
XP_055640851.1 NLVKSKYNQLPEN-----DTVTAAYKTMEPSVSGECETLYDVNAIPKY-LIQSHKDWVPQPQYMERD 61
AAQ92366.1     HTYKSKYNQFPGN-----NSFTGVYKVMEPVVTGKCKTLYDVSVVPPY-MIQANKQWVPQPQLREEG 61
Q16927.2       NLMHSSKPIHPSK-----NEWNGHYKVMEPLVTGECETHYDVNLIPAY-MIQAHKQWVPQGQLRGED 61
XP_062553809.1 NRMHSSQPVHPSK-----GEWNGHYKVMEPLVTGECETHYDVNLIPAY-MIQSHEYWVPRHQLREED 61
XP_062553910.1 NRMHSSRPVHPNK-----GEVNCHYKVMEPLVTGECETHYDVNLIPAY-MIQAHKQWVPQGHLRGED 61
AAV31928.1     HMVESKYNQFPGN-----NSFTGVYEVMEPVVTGECKTLYDVNVIPPY-IVQANKHWVPQPQLREND 61
AAA99486.1     NLMHSSKPIHPSK-----NEWNGHYKVMEPLVTGECETHYDVNLIPAY-MIQAHKQWVPQGQLRGED 61
XP_001660818.2 NLMHSSKPIHPSK-----NEWNGHYKVMEPLVTGECETHYDVNLIPAY-MIQAHKQWVPQGQLRGED 61
XP_019542945.3 HMYKSKYNQVPGN-----NSFTGMYEVMEPVVTGKCKTLYDVSVIPPY-LVQANKHWVPLPQLREEG 61
AAA18221.1     NLMHSSKPIHPSK-----NEWNGHYKVMEPLVTGECETHYDVNLIPAY-MIQAHKQWVPQGQLRGED 61
KXJ77091.1     HMYKSKYNQVPGN-----NSFTGMYEVMEPVVTGKCKTLYDVSVIPPY-LVQANKHWVPLPQLREEG 61
XP_019542959.3 NLMHSSKPVHPSK-----DEWNGHYKVMEPLVTGECETHYDVNLVPAY-MVQAHKYWYPQGQFRGED 61
KXJ71699.1     NLMHKSKPIHPSK-----DEWNGHYKVMEPLVTGECETHYDVNLVPAY-LIQAHKEWVPLEHMRGED 61
AAQ92367.1     NLMHSSKPVHPSK-----NEWNGHYKVMEPLVTGECETHYDVNLVPAY-MIQAHKQWVPQGQLRSQD 61
XP_019542943.2 NLMHKSKPIHPTK-----DEWNGHYKVMEPLVTGECETHYDVNLVPAY-LIQAHKEWVPLEHMRGED 61
XP_055857779.1 NVITCRHNQLPSD--SSTSATNAFYKTMEPTVTGECETLYEMSAVPAFSDAEPAEKWAPQTGLSMDE 65
EAT42291.1     HTYKSKYNQFPGN-----NSFTGVYEVMEPVVTGKCKTLYDVSVVPPY-MIQANKQWVPQPQLREEG 61
XP_055857778.1 NVITCRHNQLPSD--SSTGATNAFYKTMEPTVTGECETLYEMSAVPAFSDAEPAEKWAPQTGLSMDE 65
AAV31925.1     NLMHKSKPIHPTK-----DEWNGHYKVMEPLVTGECETHYDVNLIPAY-LIQAHKEWVPLEQLRGED 61
AAV31927.1     NIMHSSKPAHPSK-----GEWNGYYKVMEPLVTGECETSYDVNLVPHY-LIQSHKQWVPQAQLRKDD 61
XP_055690305.1 NEIKCKYNQMPEY--DAHKSIVGAYKTMEATVTGKCEVQYRVSPVPAQ-FAETHREWFPIPELKQNH 64
XP_055714484.1 NAIKCKYNQIPDA-QNPKKTIVGAYKTMESSVTGKCEVQYRVSPVPAQ-FAETHREWFPIPELKQNH 65
KAJ6636083.1   NHIGFSDNQLPRK-----DRDNAFYKTMEPLVTGNCETVYDISPIPDY-VIQSHPELAPLPELKGAD 61
KAG4070685.1   NQIQLSENQLPEK-----YHNTAFYKTMEPLVTGNCETVYEVSAIPDY-LIQAHPEWIPLPELKGAG 61
XP_037033226.1 NQIQLSENQLPEK-----YHSTAFYKTMEPLVTGNCETVYEISAIPDY-LIQAHPEWIPVPELKGAG 61
KAJ6646139.1   NAIQSRQNQLPAS-----DRNHAFYKTMESTASGQCETYYDISPIPEY-LVKSHPEWIPLPNLKGSG 61
XP_037926124.1 NLMPSPYNQLPNV--QENDIPSGLYEVMEPSITGQCLVVYEISPVADYAVPSLHDHWAESQPVPEGM 65
XP_037036446.1 NVNSLSENQLPQA-----DSNTAFYKTAES--TGHCETIYDISPIPDY-LVRSNPEWVPLPEMIERN 59
XP_059621569.1 NEVKCKYNQVPTQ--DPKKNIVGSYKAMESTVTGKCEVQYQVTPVPEH-YAETHREWFPIPELKHNH 64
```

#### *Diptera-Brachycera* (Yp1)

β sheet

predicted peptides

```
XP_030382044 RYNLKQQQEQGQK--YGNDY--DDKEQRTSSEEDYSELAKNAK 39
KAH8421137   RYNLQQQQ-QGQK--YFSDSNEDNK-QRTSSEEDYNELAKNTN 39
AAB06041     RYNLQQQQQQGQK--YNTDSSEENKRQRSSSEENYSEQAKNAN 41
XP_004524985 RYNGQQQPINGNK-DYDYGSSQGNQG-VTSSEEDYSESWKNQK 41
XP_037938121 RYS-KQQRQPNKE--YDY-----DSKQRTSSEEDYNEAWKSQK 35
P27878       RYNGQQQPINGNK-DYDYGSSQGNQG-ATSSEEDYSESWKNQK 41
XP_017467042 RYYGQQQPINSNK-DYDDGNQSSEEDYTTSSEEDYSESWKHPK 42
XP_004530498 RYHGQQQPVRGSNMDYASGEQQQ---AATSSEEDYSESWKEQK 40
XP_011212097 RYYGQQQPVNANNQDYESNERQ----QATSSEEDYSESWKQQK 39
XP_049310044 RYNGQRQPISNNQ-DYDYGNNKDNQG-ATSSEEDYSESWKNPK 41
XP_017467040 RYYGQQQPINSNK-DYDDGNQAA-----TSSEEDYSESWKHPK 37
XP_036319979 RYYGQQQPINSNK-DYDDGNQAA-----TSSEEDYSESWKHPK 37
XP_030378958 RYNLQEQQKSSQTQEQRQQKYKDY---DYTSSEESADEWKSAR 40
XP_017467038 RYYGQQQPINSNK-DYDDGNQAA-----TSSEEDYSESWKHPK 37
NP_001291655 RYNGQRQPISNNQ-DYDYGNNKDNQG-ATSSEEDYSESWKNPK 41
AGR33807     RYNGQRQPISNNK-DYDYGSNKDNQG-ATSSEEDYSESWKNPK 41
XP_023297337 RYNMKRQQPQR----YEM-SGEKY--ERTSSEEDSSE-WKKQE 35
TMW46108     RYNGQAAQPQY----IKYQDDSNENNNAGSSEEDNNNSWKNPS 39
XP_046803011 RYNGENPEPAN----VKYED-SSENRNPSSSEEDYSTSWKSKS 38
KNC26112     RYNGEAPQRSG----VKYEDDSSEKRGPSSSEED---EWNSKS 36
KAI8118888   RYNMKRQQPQR----YEM-SGEKY--ERTSSEEDSSE-WKKQE 35
AAS75326     RYNGQAAQPQY----IKYQDDSNENNNAGSSEEDNNNSWQNPS 39
XP_037811440 RYNIKRQQPQR----YEM-SGEKY--ERTSSEEVSSE-WKKQE 35
AAS75328     RNNHKQQKLQN----YKY-SEEQTG-SRTSSEENSAE-LKNIK 36
XP_011188958 RYHGQQQPVNAN---FESDERQ----PATSSEEDYSESWKQPR 36
UPI76698     RYHGQQQPVNAN---FESDERQ----PATSSEEDYSESWKQPR 36
XP_037811447 RYNRKQQKPEN----TKY-SAEQTG-PRTSSEETSSE-WQNQK 36
KAI8118889   RYNGENPEPAN----VKYED-SSENRNPSSSEEDYSTSWKSKS 38
AAS75329     RYNNKRQQPQA----FEY-SGEKM--SRTSSEEDSNE-WQNQQ 35
KNC26108     RYNGENPEPAN----VKYED-SSENRNPSSSEEDYSTSWKSKS 38
KAI8118890   RYNGENPEPAN----VKYED-SSENRNPSSSEEDYSTSWKSKS 38
XP_023297345 RYNGENPEPAN----VKYED-SSENRNPSSSEEDYSTSWKSKS 38
TMW47569     RNNHKQQKLQN----YKY-SEEQTG-SRTSSEENSAE-LKNIK 36
XP_023297336 RYNTKRQQPQR----FDY-SGEKM--ARTSSEEDSSE-WQKQE 35
XP_037811439 RYNGDNPEPAN----VKYED-SSENRNPSSSEEDYSTSWKSRS 38
KAI8118882   RYNTKRQQPQR----FDY-SGEKM--ARTSSEEDSSE-WQKQE 35
KNC26117     RYNRKQQKPEY----KKF-SEEQTG-SRTSSEETSSE-WQNQK 36
KAI8118884   RYNRKQQKPEY----KKF-SEEQTG-SRTSSEETSSE-WQNQK 36
XP_036339359 RYYGQQKPMNNFQ---DSGEKWP----ATSSEEDYSESWKQPR 36
XP_023297331 RYNTKKQQPQK----FDY-SGEKM--SRTSSEEDSSE-WQKQE 35
XP_017479930 RYYGQQKPINNFQ---DSGEKWP----ATSSEEDYSESWKQPR 36
XP_037811434 RYNGDYPEPAN----VKYED-SSENRNPSSSEEDYSTSWKSKS 38
XP_037811460 RYNLRRQQPQQ----FEY-SGEKM--SRTSSEEDSTE-WQKQQ 35
KMQ82231     KYNAQEQQQ--------QLKSSDY---DYTSSEEAADQWKSAK 32
KAI8118883   RYNTKKQQPQK----FDY-SGEKM--SRTSSEEDSSE-WQKQE 35
AAC01961     RYYGQQQPMDINSKEYDYGSISGNK--ISSSSEEDNDSSKNPR 41
XP_023297341 RYNGEAPQRSG----VKYEDDSSEKRGPSSSEED---EWNSKS 36
CAA50066     RYSTKRQQPSK----FDY-SGEKM--ARTSSEEDSNE-WQNQQ 35
XP_054728805 RYYGQQQPIDINSKEYDYGSINGNQ--ISSSSEEDNDSSKNPR 41
XP_053947951 RYYGQQQPIDINSKEYDYGSISGNK--ISSSSEEDNDSSKNPR 41
TMW46109     RYNNKRQQPQA----YEY-SGEKM--SRTSSEEDSNEEWQNQQ 36
XP_053951037 RYYGQQQPIDINSKEYDYGSISGNK--ISSSSEEDNDSSKNPR 41
XP_023297334 RYNRKQQKPEY----KKF-SEEQTG-SRTSSEETSSE-WQNQK 36
AAS75325     RYDNKRQQPQA----YEY-SGEKM--SRTSSEENSNE-WQNQQ 35
XP_037938122 RYSGQQQQQQSD---Y-------DKNQRSSSEEDSNESQKNSN 33
XP_054727600 RYYGQQKPIDINSKEYDYGSISGNQ--ISSSSEEDNDSSKNPR 41
XP_046802539 RYNLRRQQPQQ----FEY-SGEKM--SRTSSEEESNE-WEKQQ 35
TMW46110     RYNGPAPEPQN----IKYDDESNETNNASSSEEDNAAAWKNGS 39
XP_030382019 RYNLQSYRSQ-----YQYGEQDNDDSEEQQQLQGRSKQNGDNN 38
XP_037811459 RYNGEAPQRSG----VKYEDDSSEKTGPSSSEEY---EWSSQS 36
AAS75327     RYNGPAPEPLN----VKYESASNEQINVGSSEEE----WKNGS 35
XP_061395320 RYNVQRQKPRS----YDY-SSEK-----NSERDSSSE-WETPK 32
XP_037811458 RYNGEAPKRNS----VKYEDDSSEKTSPSSSEED---ELNRNS 36
XP_023297343 RYNGEAPKRSS----VKYEDDSSEKKGSSSSEED---EVNSNS 36
KAH8421136   RYNLQFYPDQ-----YDDSDEQKSSSEVQQQ-QPR-KQNRDKD 36
P27587       AYNGRVQVQGE------QGDDSNQ---DTSSSEESSNRPNGQQ 34
UPI76699     AYNGQVQVQGG------QNDDSEQ---DTSSSEEASSRSSSQ- 33
XP_005191012 HYNVQRQKPRS----YDYDSSEKK---TSTRRDYDDE-WETPK 35
XP_018787290 AYNGQVQVQGG------QYADSEQ---DTSSSEESFN--SKQT 32
XP_004524984 AYNGRVQVQGE------QGDDSNQ---DTSSSEESSNRPNGQQ 34
XP_017467043 MYNGQIQVQIN------HGDEFEQ---AVSSSEETTS---QQR 31
XP_011191709 AYNGQVQVQGG------QNDDSEQ---DTSSSEEASSRSSSQP 34
XP_013104425 RYNRDSESSVH-----SYAAYDNEKRQRNNEEDKYSASYAKNY 38
```

*Tardigrada* (Vg)

```
                                                 β-sheet
                     predicted peptides
GAU88667.1        VLNVTRSWNYDNCKNRPIHLESLNNPQKC
GAU88362.1        VLNVTKTWDRKNCEYRPFVYNGFHLSNEC
XP_055338826.1    ILNVTKSWDYTNCNNRPIEFQSLNNPIKC
OQV21904.1        VLNVTKTFDYRNCEFQPFVFKGIHLTSTC
OQV17559.1        VLNVTKTWNYDNCRSRPITLESLNSGAKC
XP_055331266.1    VLNVTKTFDRRNCEFRPFVFKGLHLTTAC
```

*Acarina* (Vg)

β-sheet

predicted peptides

```
EEC14774.1      YVLNFTKTKNYHKCV--GKTTVFQHVDYEH 28
QBA99605.1      SVLNLTKTKNYHKCL--GYTAVFHHADYEY 28
XP_037555480.2  SVLNLTKSKNYHKCL--GHTSVFHHADYEY 28
AGQ57040.1      TVLNLTKTKNYHKCL--GHTAVFHHADYEY 28
XP_037287793.1  TVLNLTKTKNYHKCL--GLTSVFHYADYEY 28
UCJ02328.2      SVLNLTKTKNYHKCL--GYTSVFHHVDYEY 28
ABW82681.2      SVLNLTKTKNYHKCL--GRTAVFHYADYEY 28
KAK8759090.1    SVLNLTKTKNYHRCL--GHTAVFHHADYNY 28
XP_064476439.1  TMLNFTKVVNPKKCI--GKVPTFKYAAYDS 28
BAH02666.2      AVLNFTKVINHNMCLKGGKVPTFKYAAYDS 30
AGQ56699.1      QQLNLTKTRNYMKQM--GPDGRMLQNGHER 28
XP_018494463.1  QQLNLTKTRNYMKQV--GPDGRYLQNGHDG 28
OQR67440.1      NYFNVTKTRNYMQQV--GRDGRYFQNGHDE 28
QBZ96191.1      QQLNLTKTRNYMKQV--GPDGRMLQNGHDR 28
AFN88464.1      QQLNVTKTRNYMKQI--GRDGRYFQNGHDW 28
QCX36526.1      QQLNLTKTRNYMKQI--GPDGRWLQNGHER 28
```

*Cypriniformes* (Vg)

```
                              β-sheet
                              predicted peptides
NP_001038378.1  VISEDPKADRITVTKSRDLSHCQERIVKDIGLAY  34
XP_026053862.1  VISEDPKANHITVTKSKDLSHCQERIMKDVGLAY  34
XP_056125655.1  VINEDTKANHIIITKSKDLNHCQERIIRHTGLVY  34
KAK7126994.1    VISEDPKANNITITKSKDLNHCQERIIRHTGLAY  34
TRY90352.1      VINADTKVNQVYITKSKDLTNCQERIIKDIGLAY  34
XP_056125726.1  VINEDTKANHIIVTKSKDLNHCQERIKKDVGLAY  34
XP_051550339.1  VINEDPKANHIIVTKSKDLSHCKERIKKDVGLTY  34
AIR92104.1      VISEDPKANHVIVTKSKDLSHCQERIMKDVGLAY  34
XP_039524615.1  AISEDPKANHIIVTKSKDLNQCQERIMKDVGLAY  34
XP_057181767.1  IINEDLKANHIIVTKSKDLSHCQERIMKDIGLAY  34
AAD23878.1      AINEDTKANHIIVTKSKDLNQCQERIMKDVGLAY  34
XP_052445690.1  VISEDPKANHITVTKSKDLSHCQERIMKDVGLAY  34
XP_043078189.1  VINEDPKANHITVTKSKDLSHCQERIIKDVGLAY  34
KAI7791430.1    IINEDLKANHIIVTKSKDLSHCQERIMKDIGLAY  34
AIE43955.1      VINEDQKANHIIVTKSKDLSHCHDRIIKDIGLAY  34
AIE43959.1      VINEDQKANHIIVTKSKDLNHCQDRIMKDIGLAY  34
KAF4097788.1    VISEYPKANHITVTKSKDLSFCQDRIIKDVGLAY  34
KAG1928328.1    AINEDTKANHIIVTKSKDLNQCQERIMKDVGLAY  34
XP_042605319.1  VINEDPKSNHITVTKSKDLSHCQERIMKDIGLAY  34
WPD49410.1      VISEDTKANHIIVTKSKDLSHCQERIMKDIGLAY  34
KAK9967421.1    VISEDTKANHVIVTKSKDLSHCQERIMKDVGLAY  34
ROL40727.1      VISEDTKANHIIVTKSKDLSHCQERIIKDVGLAY  34
WPD49409.1      VISEDTKANHIIVTKSKDLSHCQERIMKDIGLAY  34
ATX74744.1      IISEDPKANHIIVTKSKDLSHCQERIMKDIGLAY  34
XP_016311024.1  VINEDPKTNHITVTKSKDLSHCQERIMKDVGLAY  34
XP_016112087.1  VINEDPKTNHITVTKSKDLSHCQERIMKDVGLAY  34
XP_051763597.1  VISEDPKANHIIVTKSKDLSHCQERIMKDVGLAY  34
KAK2907071.1    VISEDLKANHIIVTKSKDLRHCQERIIKDVGLAY  34
XP_051977522.1  VINEDPKANHIIVTKSKDLSHCKERIMKDIGLTY  34
```

*Atherinomorphae* (Vg)

β-sheet

predicted peptides

```
NP_001098310.1 SKADRIHLSKTKDLNHCQERIYKDVGLAGYTESCT 35
XP_061594362.1 PKADRIHLTKTKDLNHCQERIIKDIGWTGYTEKCA 35
XP_037553994.1 SKADRIHLTKTRDLNHCPDRVYLDFGLAGYIDKCV 35
XP_012721455.2 SKADRLHLTKTTDLNHCTDSIHMDVGMAGYTEKCA 35
XP_047230925.1 SKADRLHLTRTTDLNHCTDRIHMDFGMAGFTDKCA 35
XP_015234160.1 SKADRLHLTKTTDLNHCTDRIYKDVGIAGFAEKCS 35
Q98893.1       SKADRLHLTKTTDLNHCTDSIHMDVGMAGYTEKCA 35
MEQ2275361.1   SKADRLHLTKTTDLNHCTDRIHMDFGMASYTDKCA 35
XP_032429086.1 PKADRLHLTKTTDLNQCSEKIYMNVGMAGYTDKCA 35
BAD93698.1     PKADRLHLTKTTDLNLCTEKINMDVGMAGYTNKCE 35
MEQ2296058.1   SKADRLHLTKTTDLIHCTDRIHMDFGMAGYTDKCA 35
XP_007542549.1 PKAERLHLTKTTDFNHCSEKIYMDVGMAGYTEKCA 35
XP_054901662.1 PKAERLHLMKTTDLNQCSQKINRDVGMAGYAVKCE 35
XP_041861372.1 SKAERIHLTKTKDLNHCQERIFKDFGLAGYIERCA 35
NP_001316273.1 PKAERIYLTKTKDLNHCADGVSMDVGLAGYVDRCA 35
MED6284428.1   SKADRLHLTKTTDLNHCTDRIHMDFGMAGYTDKCP 35
MED6233005.1   SKADRLHLTKTTDLNHCTDRIHMDFGMAGYTDKCA 35
MEQ2236789.1   SKADRLHLTKTTDLNHCTDRIHMDFGMAGYTDKCA 35
XP_054608292.1 SKDERIHITKTRDLDQCSDKVYKDIGMAAFSDRCS 35
ACI30218.1     SKADRLHLTKTTDLIHCADRIHMDFGMAGYTDKCA 35
XP_013855644.1 SKADRIHLTKTKDLTNCPDRVYMDIGLAGYADRCV 35
CAG5864437.1   SKAERIHLTKTKDLNHCQERIFKDVGLAGYTERCS 35
```

*Decapoda* (Vg)

```
                                  β-sheet    β-sheet           β-sheet
                                        predicted peptides
BAB69831.1      SITCKDKNIIKPAYGSYKYVEAHQESVLRFQSKTD 35
AQX37249.1      SITCKDKNIIKPAYGSYKYIEAHQESVLRFQSETD 35
AJP60219.1      SITCKDKNVIKPAYGSYKYVEAHQESVLRFQSETD 35
AFM82474.1      SITCKDKNIIRPAYGSYKYIEAVQESTLRFESETD 35
BAD11098.1      TVSCKDKNIIRPAYGSYKYIEAIQESELRFQSQTD 35
UWT50543.1      SITCKDRNIIRPSFGSYKYVEAHQESTLRFHSETD 35
XP_042241566.1  SIICEDKKVVRPSYGAYKYVEAKQESTLRYVSLSS 35
AAG17936.1      AITCEDKNVVKPSYGAYKYVEAKMMSTLKYLSESS 35
BAD98732.1      SIMCHDKNIVRPAIGIYQYVEAHQESTLHFISETT 35
AAN40700.1      AITCEDKHIVRPAFGLYKYVEANQESTLRFISESS 35
UKG18870.1      KIICEDNNTIRLSYGAYRYVVATQKSSFSYLSESS 35
BAF91417.1      SIQCEDWNKVRPSYGAYKHVEARQESTLRYQSQSS 35
XP_045133432.1  SVVCEDKKVIRPSYGIYKYVEAKQESTLKLTSSDV 35
KAK7073330.1    SITCEDKHIIKPTFGSYKYVEAIQVSRLHFQSESQ 35
AKI23633.1      SVKCEDKKVVRPSYGMYKFVEAEQESTLRLVSSDA 35
XP_050717478.1  SVKCEDKKVVRPAYGTYKFVEAEQESTLRLTSSDV 35
XP_063861656.1  SVVCEDKKVIRPSYGIYKYVEAKQESTLRLTSSEA 35
AEI59132.1      SVVCEDKKVIKPSYGIYKYVEAKQESTMKLTSSDV 35
AAU93694.1      SVVCEDKKVIRPSYGMYKYVEAKQESTLRLTSSDV 35
ALE30145.1      AITCEDKNIVRPAFGLYKYVEANQESTLRFIS--- 32
```
